# Membrane-assisted assembly and selective autophagy of enteroviruses

**DOI:** 10.1101/2021.10.06.463375

**Authors:** Selma Dahmane, Adeline Kerviel, Dustin R. Morado, Kasturika Shankar, Björn Ahlman, Michael Lazarou, Nihal Altan-Bonnet, Lars-Anders Carlson

## Abstract

Enteroviruses are non-enveloped positive-sense RNA viruses that cause diverse diseases in humans. Their rapid multiplication depends on remodeling of cytoplasmic membranes for viral genome replication. It is unknown how virions assemble around these newly synthesized genomes and how they are then loaded into autophagic membranes for release through secretory autophagy. Here, we use cryo-electron tomography of infected cells to show that poliovirus assembles directly on replication membranes. Pharmacological untethering of capsids from membranes abrogates RNA encapsidation. Our data directly visualize a membrane-bound half-capsid as a prominent virion assembly intermediate. Assembly progression past this intermediate depends on the class III phosphatidylinositol 3-kinase VPS34, a key host-cell autophagy factor. On the other hand, the canonical autophagy initiator ULK1 is shown to restrict virion production since its inhibition leads to increased accumulation of virions in vast intracellular arrays, followed by an increased vesicular release at later time points. Finally, we identify multiple layers of selectivity in virus-induced autophagy, with a strong selection for RNA-loaded virions over empty capsids and the segregation of virions from other types of autophagosome contents. These findings provide an integrated structural framework for multiple stages of the poliovirus life cycle.

## Main

Enteroviruses are a major genus of positive-sense RNA viruses within the Picornaviridae family. They cause a wide variety of human diseases such as poliomyelitis (poliovirus), related acute flaccid myelitis conditions (e.g. EV-D68), and viral myocarditis (Coxsackievirus B3). The enterovirus particle is a non-enveloped particle of ∼30 nm diameter, encapsidating a single-stranded RNA genome of about 7500 nucleotides.

Upon infection, enteroviruses rapidly remodel cytoplasmic membranes to create an optimal environment for virus replication^1-3^. This starts with disassembly of the secretory pathway, in particular the Golgi apparatus as seen by dispersion of Golgi marker proteins throughout the cytoplasm^4^. Ensuing single membrane tubules and vesicles are the site of viral RNA replication, but little is known about the site of virion assembly^3,5^. The viral genome has preferential interaction sites with the capsid but contains no known high-affinity packaging signal sufficient for RNA loading into capsids^6,7^. This suggests that virion assembly may take place in immediate vicinity of RNA production sites to ensure specific RNA encapsidation, which is supported by the finding that the viral membrane-bound helicase 2C interacts with the capsid protein VP3^7,8^.

A biochemically distinct late stage of replication starts around 6 hours post-infection (h p.i.). It is characterized by the lipidation of host proteins belonging to the LC3 subfamily of ATG8s, a hallmark of the autophagy pathway^9^. Induction of autophagy in infected cells is mediated by the viral protein 2BC and is independent of the ULK1/ULK2 protein kinases that initiate canonical autophagy^9,10^. At this late stage, double-membrane structures reminiscent of autophagic membranes are formed in the infected cell^11^. The inherently non-enveloped picornaviruses have recently been shown to leave cells non-lytically as groups of virions contained in LC3-positive lipid vesicles^12-16^. It thus seems plausible that autophagy-like double-membrane structures observed in infected cells relate to non-lytic virus egress. However, conventional EM sample preparation is too destructive to macromolecular structure to allow an *in situ* analysis of autophagosome contents - the experiment needed to directly test this hypothesis.

To shed light on how enteroviruses are assembled and packaged into autophagosomes, we took advantage of recent advances in focused-ion-beam milling and cryo-electron tomography^17-21^. The *in situ* structures of poliovirus-infected cells revealed that enteroviruses assemble directly on replication membranes. Completion of virus assembly requires the host lipid kinase VPS34, and RNA loading into virions correlates with their membrane tethering. Inhibiting the initiation of canonical autophagy surprisingly increased virion production and release. The cryo-electron tomograms further revealed that virus-induced autophagy has a striking degree of selectivity, selecting RNA-containing virions over empty capsids, and segregating virions from other types of autophagosome contents.

## Results

### Cryo-electron tomography reveals that poliovirus RNA loading requires capsid tethering to membranes

To investigate enterovirus assembly *in situ*, we infected HeLa cells with poliovirus type 1 and imaged the cytoplasm with cryo-electron tomography at different time points post infection (Supplementary fig. 1). At 3 h p.i. first new virions had already assembled in the cells (Supplementary Fig. 2A-D). Tomograms recorded at 6 h p.i. showed a more starkly remodeled cytoplasm (Fig. 1A-B). Compared to uninfected cells and 3 h p.i., there was a significant increase in both open cup-shaped structures resembling phagophores and closed double-membrane vesicles (Fig. 1A-B, Supplementary fig. 2A-G). These membranes will collectively be referred to as autophagy-like membranes (ALMs).

**Figure 1:**
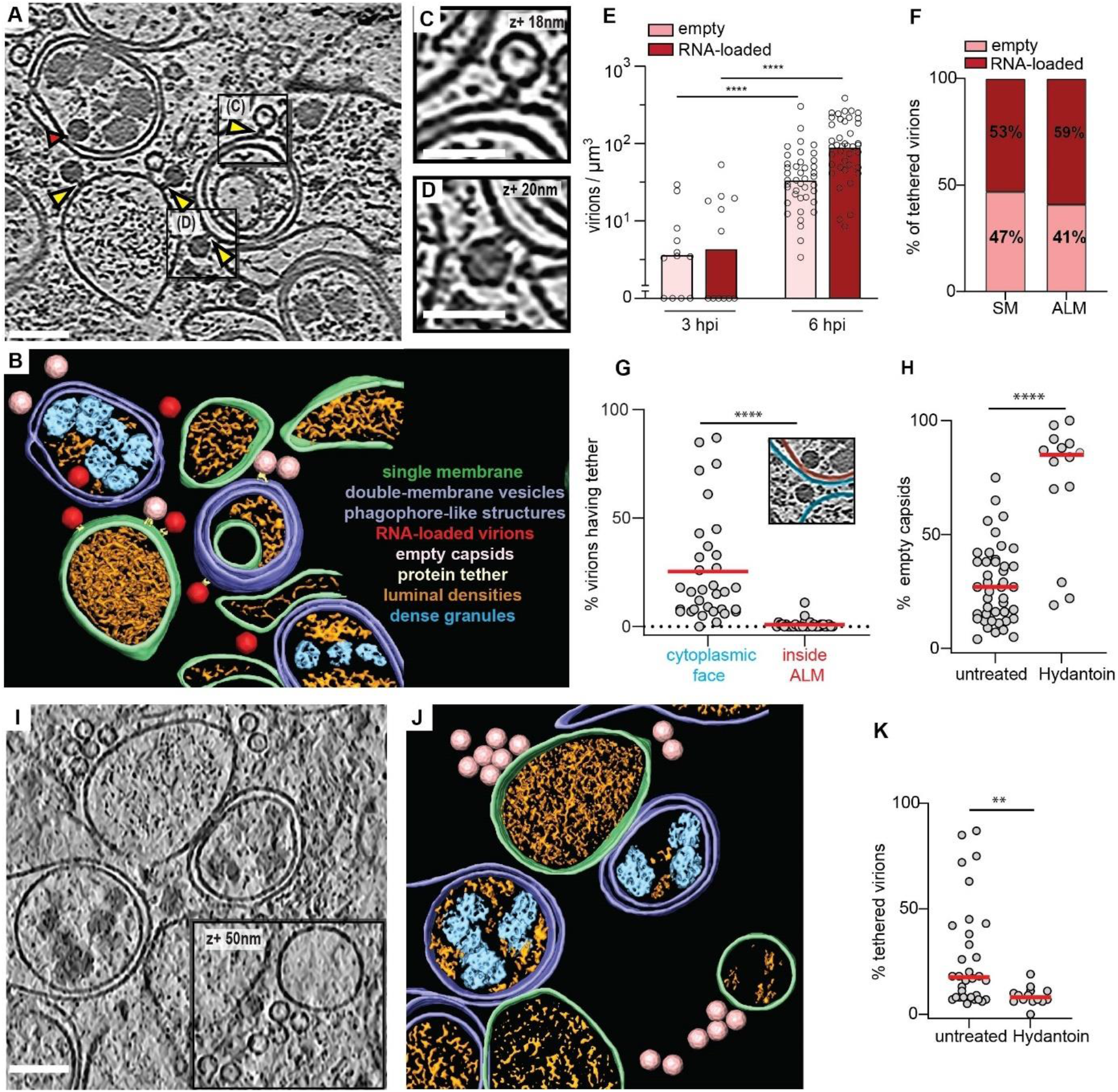
Cryo-electron tomography allows the visualization of poliovirus replication and assembly sites *in situ*. (A) Slice through a representative cryo-electron tomogram of a lamella milled through a PV-infected cell at 6 h p.i., revealing PV-induced SM (Single Membranes) and ALM (Autophagy-Like Membranes) proliferation. Yellow arrowheads indicate densities tethering intracellular empty capsids and darker RNA-loaded virions to membranes. Red arrowhead indicates a virion enclosed inside a DMV (Double-Membrane Vesicle), proximal but not tethered to the membrane. (B) Segmentation of the tomogram presented in (A). Color labels are defined for each structure. Empty capsids (pink) and RNA-loaded virions (red) are represented by their subtomogram averages. (C-D) Magnified view of an empty capsid and RNA-loaded virion tethered to a DMV shown in (A) with black boxes. (E) Concentration of intracellular empty capsids and RNA-loaded virions in tomograms at 3 h p.i. and 6 h p.i., as measured by template matching. (F) Percentage of empty and RNA-loaded virions on SM and ALM. (G) Percentage of virions on the outside (blue) and inside (red) of ALM having a visible tether to a membrane as indicated in the inset. (H) Percentage of empty capsids in tomograms of untreated and Hydantoin-treated cells at 6 h p.i., as measured by template matching. Horizontal lines represent the average. (I) Cryo-electron tomogram of a PV-infected, Hydantoin-treated cell at 6 h p.i., containing several empty capsids which are not tethered to the surrounding membranes. (J) Segmentation of the tomogram in (I). Color labels for each structure are the same in (B) and empty capsids are represented by their subtomogram average. (K) Percentage of tethered virions in untreated and Hydantoin-treated cells as observed in 6 h p.i. cryo-tomograms. Horizontal lines represent the average. In all graphs, each dot corresponds to one tomogram analyzed (see also Supplementary table 2). Statistical significance by unpaired two-tailed Student’s t test; **p<0,01 and ****p <0,0001. Scale bars: (A-I) 100 nm, (C-D) 50 nm.

From the tomograms we could clearly distinguish empty capsids from RNA-loaded virions (Fig. 1A-B). At 6 h p.i. the cytoplasmic concentration of empty and RNA-loaded particles was on average 7 and 20 times higher than at 3 h p.i., respectively, as measured by a template matching procedure (Fig. 1E). Strikingly, both empty capsids and RNA-loaded virions were frequently tethered to membranes through macromolecular complexes (Fig. 1A-D, yellow arrowheads). The tether appeared to have a defined size and keep the virions at a defined distance from the membrane. At 6 h p.i., virions were tethered to both single-membrane tubes and vesicles (SMs) and ALMs, and the fraction of empty vs. RNA-loaded virions was similar on both types of membranes (Fig. 1F). We noticed that virions were only tethered to the outer face of ALMs and SMs, whereas virions engulfed by ALMs had lost the tether (Fig. 1A, red arrow and Fig. 1G).

The visualization of capsids tethered to replication membranes suggested that capsid RNA loading took place on membranes rather than in the cytosol. To test this we used (5-3,4-dichlorophenyl)-methylhydantoin (henceforth Hydantoin), an antiviral drug that inhibits RNA loading of PV capsids^22^. Cells were infected with PV and treated with Hydantoin at a concentration of 50 μg/ml that did not interfere with viral RNA replication^22^ before processing for cryo-ET at 6 h p.i. This revealed both an increase in the fraction of empty capsids to ∼70% and a ∼3-fold decrease in the fraction of tethered capsids in Hydantoin-treated cells (Fig. 1H-K). Notably, that the abundance of SMs and ALMs remained unchanged (Supplementary fig. 2H). Together, these data show that newly assembled poliovirus capsids are tethered to the cytoplasmic face of SMs and ALMs and tethering facilitates viral RNA encapsidation.

### Enterovirus capsid assembly takes place on membranes and requires VPS34 activity

Tomograms of infected cells at 6 h p.i. frequently contained novel structures that had a size and shape consistent with partial capsids (Fig. 2A-E, Supplementary Fig. 3A-B). Strikingly, in 15 tomograms of infected cells, 96% of these *bona fide* capsid intermediates were membrane-associated (58% SM, 38% ALM) whereas only 4% were found in the cytoplasm (Fig. 2F). The capsid intermediates contained variable luminal densities and were observed at different angles to the membranes (Fig. 2B-E). Due to this structural variability, we characterized them by a simple angle of closure, which resulted in a unimodal distribution with an average of 169°, i.e. closely corresponding to half a capsid (Fig. 2G). The clear clustering around a single value indicates that the membrane-bound capsid intermediate is a single, or a set of closely related, molecular species.

**Figure 2:**
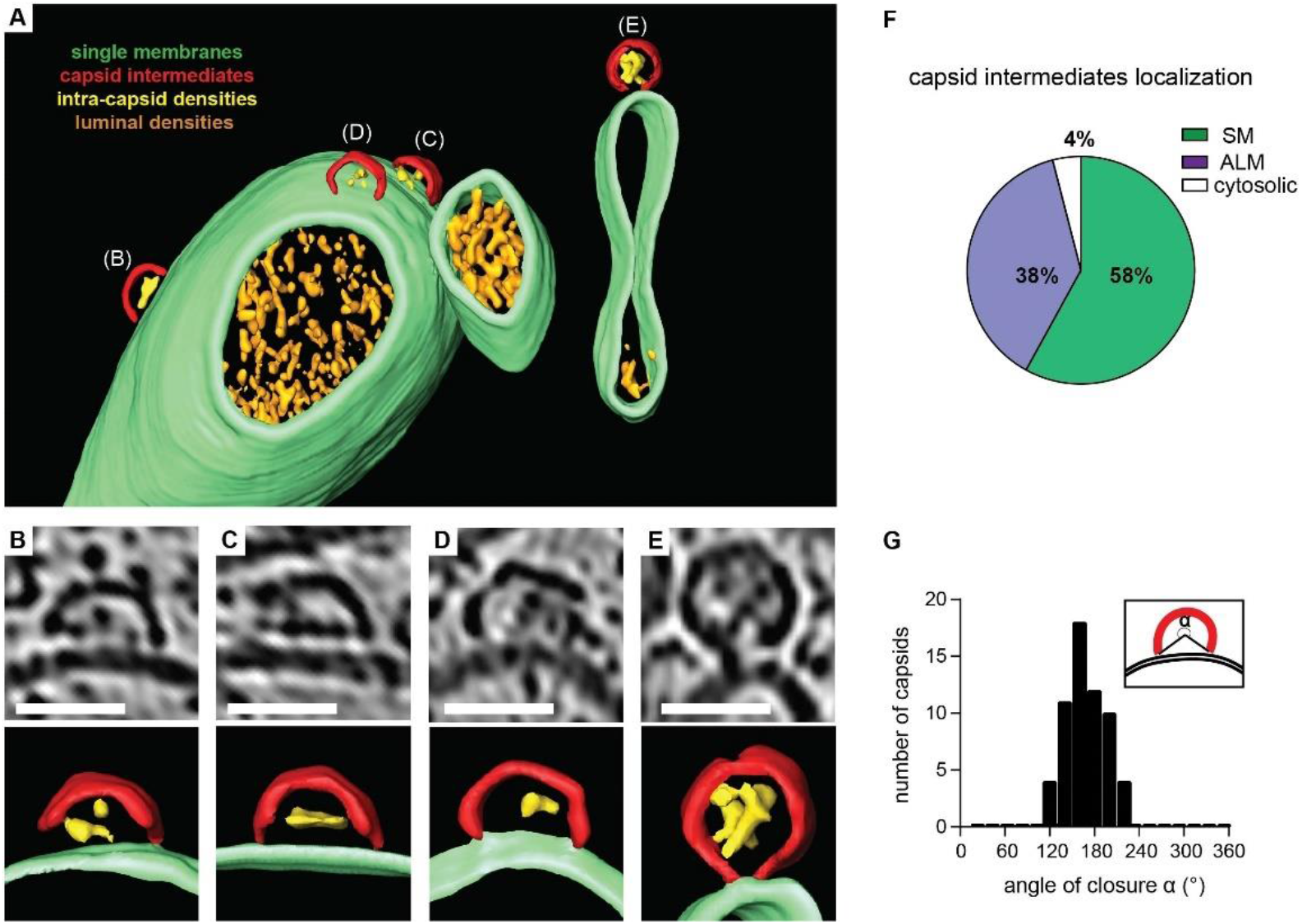
A membrane-bound capsid intermediate. (A) 3D segmentation of a tomogram showing capsid assembly intermediates, containing luminal densities, directly bound to single-membrane tubules (corresponding slice in Supplementary Fig. 3A-B). (B-E) Zoomed tomogram slices and segmentations of the capsids assembly intermediates marked in (A). (F) Percentage of capsid assembly intermediates found on SMs, ALMs or not associated with membranes, as counted in 15 tomograms at 6 h p.i. (G) Distribution of capsid intermediate closures (α), as defined in the inset, measured at 6 h p.i. Average closure was 169° (SD=26°, N=51). Scale bars: 50 nm.

Given the frequent association of partial and complete capsids with ALMs, we inhibited autophagy using two selective inhibitors: MRT68921 and Vps34-IN1. MRT68921 inhibits the ULK1/ULK2 protein kinases that initiate canonical autophagy^23^, whereas Vps34-IN1 inhibits the lipid kinase VPS34 that produces phosphatidylinositol-3-phosphate (PI(3)P) on the growing phagophore membrane^24^. Consistent with previous studies, PV induced ULK-independent LC3 lipidation at 6 h p.i.^10^. However, co-treating infected cells with both inhibitors strongly decreased LC3 lipidation (Supplementary fig. S4A). Tomograms of co-treated cells revealed reduced membrane proliferation and a 2 orders of magnitude decrease in both empty and RNA-loaded capsids (Fig. 3A-E; Supplementary fig. 4B). However, the number and distribution of membrane-bound capsid intermediates were unaffected (Fig. 3E; Supplementary fig. 4C-D). The stalled progress from capsid intermediates to full capsids meant that we could observe a rare, more advanced assembly stage that may represent the transition from the half-capsid intermediate to a complete membrane-tethered virion (Fig. 3D). Moreover Vps34-IN1 treatment alone had the same effect as the combination of inhibitors (Supplementary Fig. 4E-G). The decrease in intracellular assembled virus was mirrored by a decrease in virus release from VPS34-inhibited cells (Fig. 3F). Notably, as viral RNA replication was unaffected by inhibiting VPS34, this suggested that VPS34 activity is essential for virus assembly rather than RNA replication (Supplementary fig. 4G).

**Figure 3:**
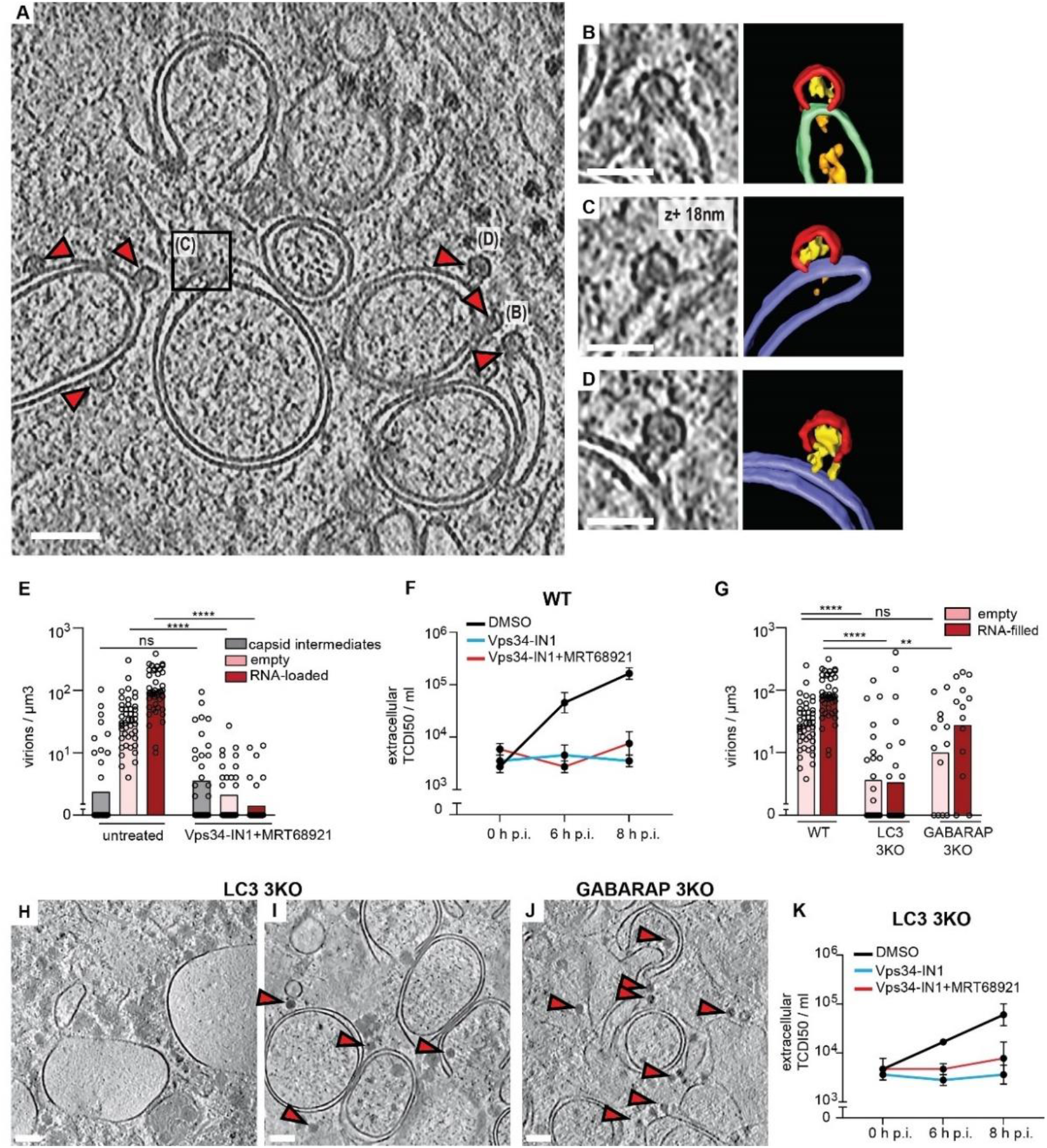
VPS34 inhibition stalls PV assembly at the half-capsid intermediate. (A) Cryo-electron tomogram of a PV-infected cell co-treated with Vps34-IN1 and MRT68921 at 6 h p.i. (B-D) Magnified views of capsid intermediates indicated in (A), and their corresponding segmentations. Capsid intermediates (red), intra-capsid densities (yellow), ALM (purple), SM (green), luminal densities (orange). (D) The magnified view shows a near-complete capsid tethered to a double-membrane vesicle. (E) Concentration of intracellular capsid intermediates, empty capsids and RNA-loaded virions in untreated and Vps34-IN1 + MRT68921 treated cells at 6 h p.i. Each dot corresponds to one tomogram, bars represent the averages (see also Supplementary table 2). (F) Released virus titer at 0, 6, and 8 h p.i. in WT cells treated with DMSO, Vps34-IN1, and Vps34-IN1 + MRT68921. Bars represent the means of biological triplicates ± SEM. (G) Concentration of intracellular empty capsids and RNA-loaded virions in in WT, LC3 and GABARAP 3KO cells at 6 h p.i. (H-I) Slices through representative cryo-electron tomograms of lamellas milled through PV-infected LC3 3KO cells at 6 h p.i., revealing two types of membrane proliferation: large single-membrane vesicles (H) and ALM proliferation (I). (J) Slice of cryo-tomogram of a GABARAP 3KO PV-infected cell at 6 h p.i., where ALMs were observed. (H-J) Red arrowheads indicate the presence of RNA-loaded virions. (K) Time course of PV release from LC3 3KO cells in the presence or absence of autophagy inhibitors as indicated in the figure. Error bars represent the means of biological triplicates ± SEM. Statistical significance by unpaired two-tailed Student’s t test; *p<0,05; **p<0,01 and ****p <0,0001. Scale bars: (A) 100 nm, (B-D) 50 nm, (H-J) 200 nm.

Since VPS34 inhibition decreased LC3 lipidation, we sought to determine if Vps34-IN1’s effect on virus assembly was mediated by ATG8 family proteins. We infected CRISPR-generated triple knock out (3KO) cells of the LC3 subfamily, and 3KO cells of the GABARAP subfamily and imaged them at 6 h p.i. The only significant change to membrane structures was a decrease in ALMs in infected LC3 3KO cells (Supplementary fig 4I). This was paralleled by a decrease of intracellular virions in LC3 KO cells (Fig. 3G). Interestingly, those areas of LC3 3KO cytoplasm that still contained ALMs also contained virions, whereas areas with large SMs were devoid of virions (Fig. 3H-I). By comparison, GABARAP 3KO cells still robustly accumulated ALMs (Fig. 3J), and there was less change in intracellular virus concentration upon GABARAP deletion (Fig. 3G). Together this indicates that PI(3)P-decorated remodeled membranes, rather than any individual ATG8 protein, are necessary for enterovirus assembly. This notion was strengthened by the observation that VPS34 inhibition further reduced virus release from LC3 3KO cells, similar to its effect on wildtype cells (Fig. 3K).

In summary, enterovirus capsid assembly takes place on membranes with a prominent half-capsid intermediate, and activity of the lipid kinase VPS34 is required for assembly to progress beyond this intermediate.

### Inhibition of ULK1 leads to formation of intracellular virus arrays and increased vesicular release

Although Inhibition of ULK1 and its homologue ULK2, the initiators of canonical autophagy, did not affect PV-induced LC3 lipidation or the generation of ALMs (Fig. 4A-B, Supplementary Fig. 5A), we found that ULK-inhibited cells contained large cytoplasmic arrays of virions (Fig. 4A-B). The arrays contained several hundred virions, virtually all RNA-loaded, in what seemed to be a close-packing arrangement. Virus arrays were also visible in freeze-substituted sections, which allowed sampling of a larger number of cells (Supplementary fig. 5B-E). Arrays were found in 3% of untreated cells and 56% of ULK-inhibited cells (Fig. 4C), a significantly higher fraction, thus indicating that ULK inhibition upregulates intracellular virus array formation.

**Figure 4:**
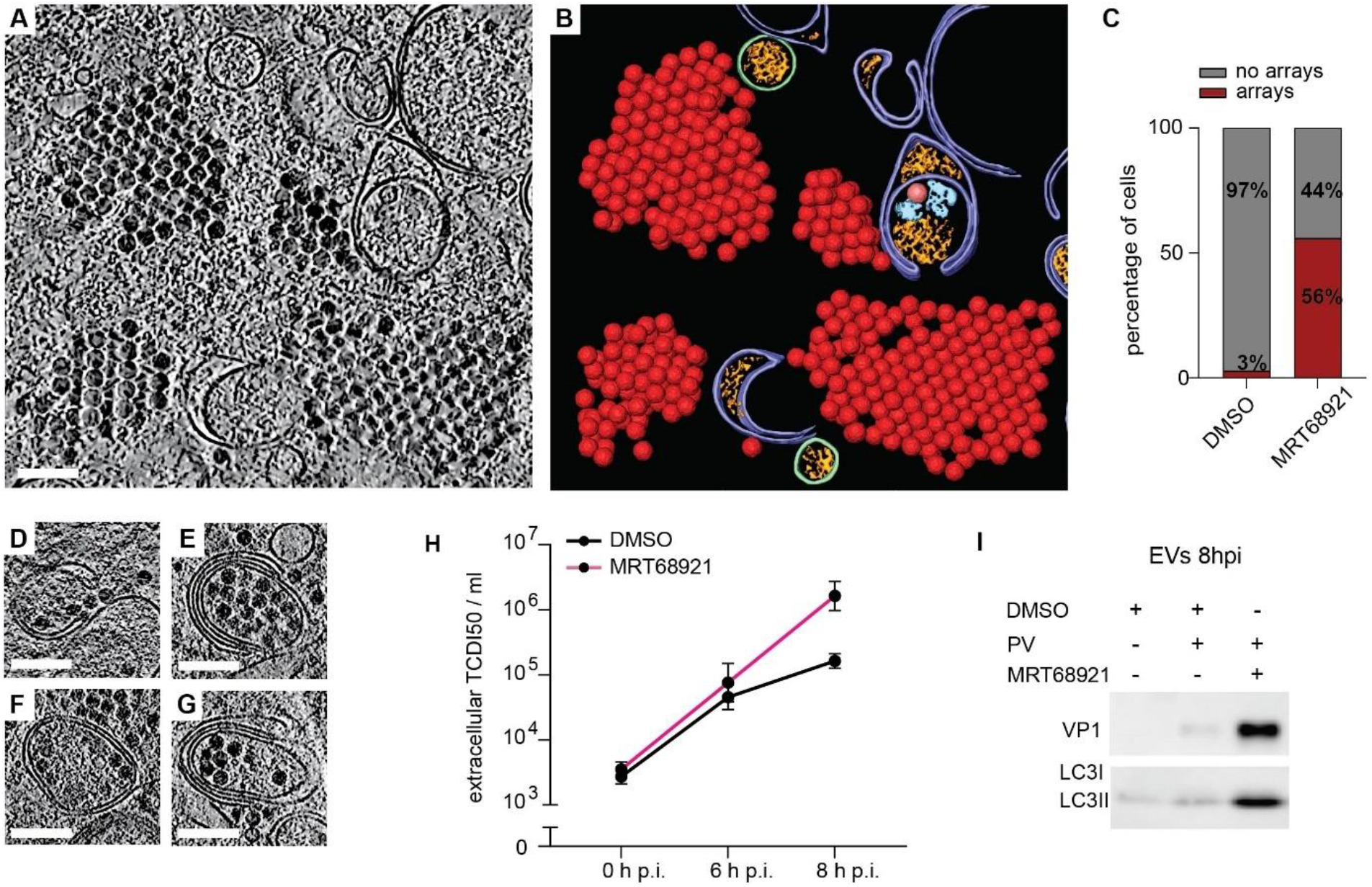
Inhibition of ULK1/ULK2 induces formation of intracellular virus arrays and increases infectious virus release. (A) Cryo-electron tomogram of an MRT68921-treated, PV-infected cell containing cytoplasmic virion arrays. (B) Segmentation of the tomogram in (A). Virions are represented by their subtomogram averages. Colors are as in Fig. 1B. (C) Percentage of cells containing virus arrays in DMSO and MRT68921-treated cells as measured from freeze-substituted sections. (D-G) Several examples of virion recruitment to ALMs in tomigrams of MRT68921-treated, PV-infected cells. (H) Time course of PV release from MRT68921 and DMSO-treated cells. Error bars represent the means of biological triplicates ± SEM. (I) Western blot analysis of extracellular vesicles (EVs) harvested at 8 h p.i. from infected cells treated or not with MRT68921. Scale bars 100nm.

The formation of virus arrays may either be due to a defect in virion release from ULK-inhibited cells, or the arrays may be part of an increased intracellular virion pool *en route* to release. The ‘release scenario’ was supported by the observation that RNA-loaded virions were abundant in ALMs in ULK-inhibited cells (Fig. 4D-G). To further discriminate between these two scenarios, we measured released infectious virus from ULK-inhibited cells at different time points and compared it to untreated cells. While extracellular PV titers were unchanged at 6 h p.i., we measured one order of magnitude increase over untreated cells at 8 h p.i. (Figure 4H). To determine if the increased virus release at late time point still took place through secretory autophagy, we isolated the extracellular vesicular fraction. A strong increase in capsid protein as well as lipidated LC3 from ULK-inhibited cells confirmed that the virions were released in vesicles positive for LC3 (Fig. 4I). Taken together, ULK activity in infected cells is not necessary for downstream autophagy processes such as LC3 lipidation but appears to put a break on the intracellular accumulation and vesicular release of virions.

### Selective packaging of RNA-loaded virion and contents segregation in autophagic membranes

The observation that virions are tethered to the outside of ALMs but not to the inside (Fig. 1G) indicated that cryo-ET can provide structural insights into the process of virion packaging into autophagic membranes. Indeed, the tomograms of PV-infected cells at 6 h p.i. showed several stages of autophagic engulfment of virions, ranging from wide open phagophores to completely sealed DMVs (Fig. 5A-E). We then re-evaluated the statistics of empty capsids and RNA-loaded virions at 6 h p.i. taking particle location (outside or inside ALMs) into consideration. Strikingly, this revealed that autophagic engulfment strongly selects for RNA-loaded virions. Outside ALMs, empty capsids represent 23% of particles, whereas only 1% of particles in ALMs were empty capsids (Fig. 5F).

**Figure 5:**
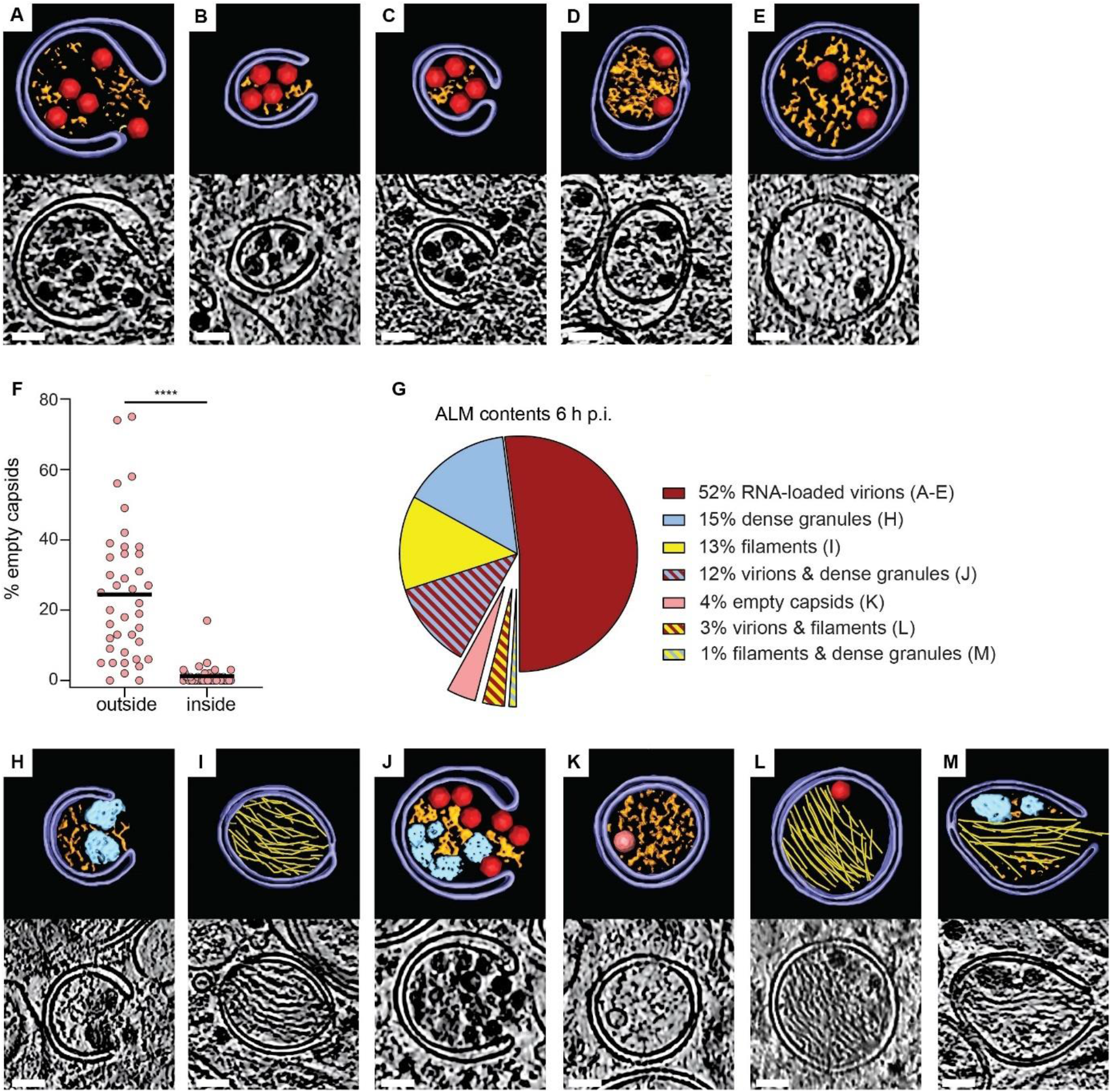
Autophagic membranes select and sort their contents in PV-infected cells. (A-E) Tomograms of PV-infected cells at 6 h p.i. showing different stages of engulfment of RNA-loaded virions by ALMs, including initial recruitment to phagophores (A-C) and enclosure in DMVs (D-E). Each panel contains a slice through the tomogram and the corresponding segmentation, colored as in Fig. 1B. (F) Percentage of empty capsids on the outside and inside of ALMs. Each dot corresponds to one tomogram analyzed; horizontal line is the average (see also Supplementary table 2). (G) Relative abundance of seven classes of ALMs by contents (single/mixed), in tomograms of PV-infected cells at 6 h p.i. (H-M) Segmentations and corresponding tomographic slices of examples of the different ALM classes, as labeled in (G). Colors are as in Fig. 1B. Protein filament bundles are shown in yellow. Statistical significance by unpaired two-tailed Student’s t test; ****p <0,0001. Scale bars 50 nm.

The tomograms allowed further classification of ALMs based on their contents (Fig. 5G-M). At 52%, the most abundant class was ALMs containing RNA-loaded virions (Fig. 5A-E,G). The second most abundant class, at 15%, was ALMs containing amorphous granular material that was clearly denser than the remaining cytoplasmic contents (Fig. 5G,H). These dense granules were frequently co-packed with virions (Figure 5G,J). One distinct class representing 13% of ALMs contained tightly packed bundles of protein filaments (Fig. 5G,I, Supplementary Fig 6A-B). As opposed to the dense granules, filaments were rarely co-packaged with other ALM contents (Fig. 5G). The filaments were not affected by VPS34 inhibition, but instead completely absent in both LC3 and GABARAP 3KO cells (Supplementary Fig. 6C). This is the opposite dependence on autophagy host factors than that displayed by virus assembly (Fig. 3G). We determined the filament structure by subtomogram averaging to 18.5 Å resolution, which yielded a helix with an average diameter of 10 nm, 29° twist and 52 Å rise per subunit (Supplementary Fig. 6D-E). From this map the identity of the filament could not be positively determined, but the map did allow definitive exclusion of protein filaments with known structure. A systematic comparison of the filament to all relevant mammalian protein filaments in the electron microscopy database allowed exclusion of all except decorated actin filaments (Supplementary Fig. S7 and Table S3 for complete list). Thus, the filament is either actin decorated by a vinculin-like actin-binding protein, or an unknown viral or cellular protein filament of similar size and shape.

Altogether, these data reveal an exquisite specificity in virus-induced autophagy: autophagic membranes selectively engulf RNA-loaded virions while excluding empty capsids, and co-package virions with dense granular material while segregating them from a second class of ALMs that contains bundles of protein filaments.

## Discussion

Here we present an *in situ* structural analysis of enterovirus replication by cryo-electron tomography. Our study focuses on the involvement of autophagic membranes in virion assembly and egress, and the data are consistent with the model presented in Fig. 6. Compared to viral RNA replication, much less is known about the site and pathway of enterovirus assembly. A membrane-proximal location of the assembly has been suggested, but direct evidence has been missing. In tomograms of infected cells, we identified an abundant capsid assembly intermediate structurally equivalent to half of an enterovirus capsid (Fig. 2). It was to 96% found directly docked to membranes in infected cells, and at roughly equal proportions on the cytosolic face of SMs and ALMs, which suggests that these two membrane types both serve as virus assembly platforms. Our findings are consistent with ultracentrifugation studies of enterovirus-infected cells that detected an abundant capsid-related species equivalent to half of an empty capsid, without being able to elucidate its identity^25,26^.

**Figure 6:**
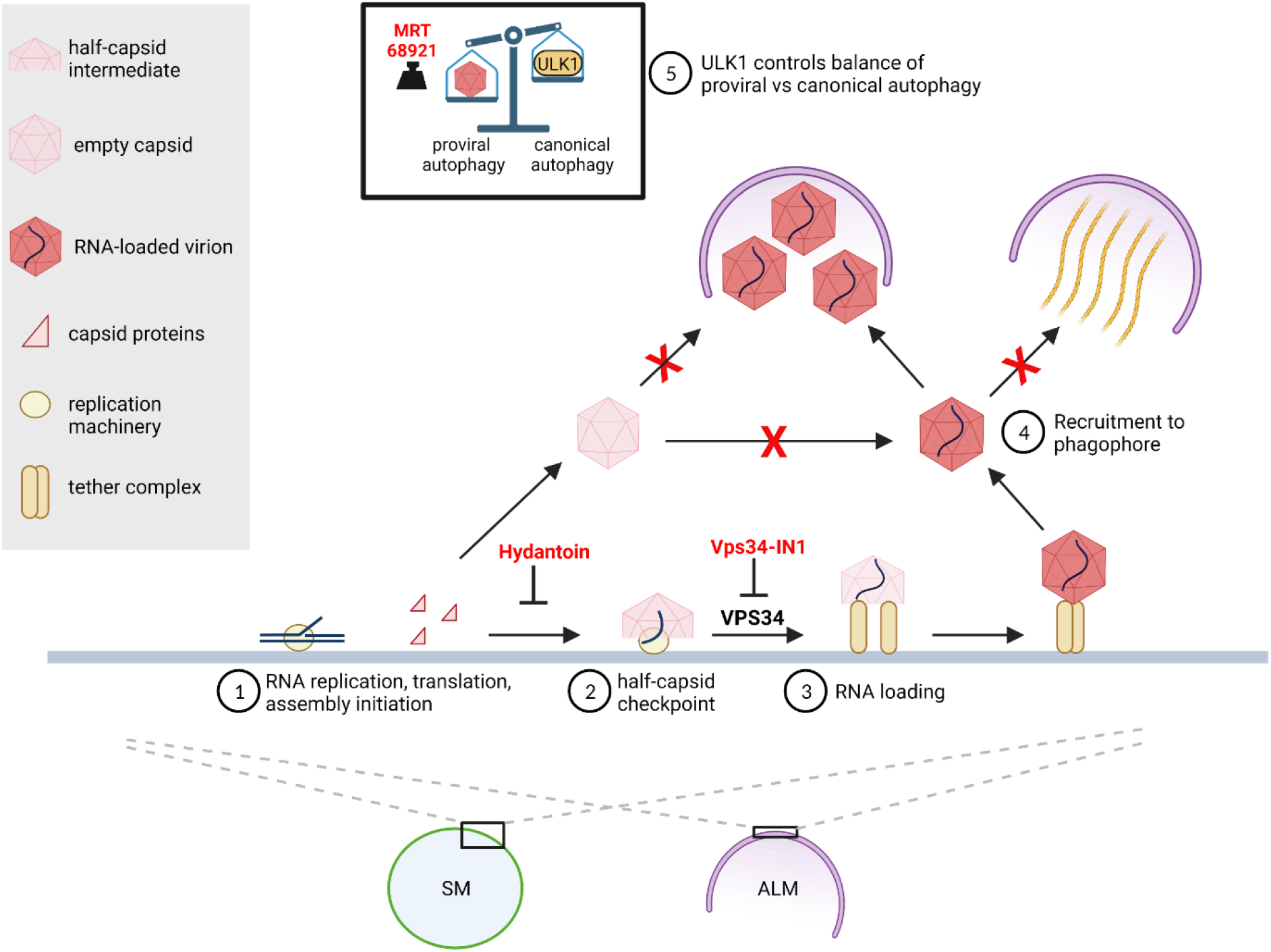
The interplay between autophagy and enterovirus replication is dynamic, selective and sequential. This model illustrates how remodeled cytoplasmic membranes act as a platform for viral RNA replication and as a production line for virion assembly. 1) Capsid proteins, produced as part of the membrane-associated viral polyprotein, assemble on the replication membrane to the point of half-capsid intermediate. The antiviral drug Hydantoin leads to premature capsid release from membranes. Away from the replication membrane, capsids cannot be loaded with RNA. 2) The assembly pauses at the long-lived half-capsid intermediate that carries out the “late proofreading” resulting in exclusive incorporation of newly synthesized membrane-associated viral RNA^28^. The inhibition of VPS34 stalls the assembly at the half-capsid checkpoint. 3) RNA-loading of the capsid leads to the formation of complete virions tethered to the replication membrane. 4) Phagophores selectively package RNA-loaded virions while excluding empty capsids. The virion-containing class of phagophores is distinct from a second class of phagophores containing bundles of protein filaments. 5) A balance between ULK1-induced canonical autophagy and virus-induced proviral autophagy regulates the level of virion production. Inhibition of remaining ULK1 activity by MRT68921 further increases virion production.

The current paradigm of non-enveloped virus assembly holds that the proteinaceous capsid either assembles independently followed by energy-expending genome loading, or is templated by the genome^27^. Our finding extends that paradigm with a third mode: assembly that is assisted by the replication membrane and its bound components. A slow transition past the half-capsid intermediate may help ensure that the viral RNA has been incorporated before the capsid closes. This is consistent with a previously proposed “late proofreading” mechanism that aimed to explain why only recently produced viral RNA, present on the replication membrane, is incorporated into virions^28^.

We showed that the lipid kinase VPS34 is a host factor required for assembly to progress past the half-capsid intermediate (Fig. 3). The VPS34 requirement may explain why the membrane-bound picornaviral helicase 2C binds the PI3 kinase complex^29^, in which VPS34 is the catalytic subunit. Hence, 2C may enable the virus to activate VPS34 independently of ULK1-dependent canonical autophagy induction. In fact, we show that pharmacological inhibition of ULK1 further boosts virus assembly and release (Fig. 4), and it was previously reported that poliovirus infection partially depletes ULK1^10^. The picture that emerges is that the virus optimizes the cellular environment by suppressing the master switch of canonical autophagy, ULK1, while at the same time activating necessary subsystems of the autophagy pathway, such as VPS34, in alternative ways.

The cryo-electron tomograms revealed several layers of selectivity in enterovirus-induced autophagy (Fig. 5). Perhaps most remarkably, there is a strong selectivity for packaging of RNA-loaded virions over empty capsids. Selectivity had not previously been demonstrated for virus-induced autophagy, but it is frequent in other forms of autophagy. There, so-called autophagy receptors mediate the degradation of specific cytoplasmic components by linking them to LC3 on the growing phagophore membrane^30^. The case of enterovirus particles thus seems to present a conundrum: How can phagophores selectively package RNA-loaded virions over empty capsids when the external surfaces of both these particles are virtually identical^31^? Further studies will be needed to elucidate this mechanism, but a clue may be provided by the correlation between RNA loading of capsids and their membrane tethering, as well as the loss of tethering upon autophagic engulfment (Fig. 1).

The tomograms allowed further structural catalogization of ALM contents in infected cells (Fig. 5). Viruses were frequently copackaged with dense granular material which has also been seen in tomograms of vesicles released from enterovirus-infected cells^32^. The high electron density of the granules is compatible with them containing RNA, which could mean that non-encapsidated viral or cellular RNA is released in the same vesicles as virions. On the other hand, virions were markedly segregated from ALMs that contained bundles of protein filaments. The filaments had a structural signature compatible with decorated F-actin. This tentative identity of the filaments would tie together previous reports that the actin cytoskeleton largely disappears in both starved and enterovirus-infected cells^33,34^. In starved cells this is linked to presence of F-actin in the lumen of LC3-positive membranes (where it was suggested to play a role in shaping the phagophore)^34^. It is thus possible that the filament-filled ALMs represent structural snapshots of this process.

In summary, our study of poliovirus-infected cells by cryo-electron tomography reveals membrane-assisted capsid assembly and a link between membrane tethering and RNA-loading of virions. It further shows the multi-faceted nature of autophagy in enterovirus-infected cells, balancing virion production and autophagic engulfment, making sure that only RNA-loaded virions are packaged in phagophores and segregating them from other types of autophagic cargoes.

## Acknowledgments

We would like to thank Carlson lab members, Sven Carlsson, and Richard Lundmark for constructive suggestions, Dale Corkery for help with the LC3 lipidation assay, as well as Nitesh Mistry for help with virus purification (all Umeå University). We are also thankful to Sebastian Schultz and Andreas Brech (Oslo University) for sharing their experience in freeze substitution.

Cryo-EM data were collected at the Umeå Centre for Electron Microscopy (UCEM), a SciLifeLab National Cryo-EM facility and part of National Microscopy Infrastructure, NMI VR-RFI 2016-00968), supported by instrumentation grants from the Knut and Alice Wallenberg Foundation and the Kempe Foundations.

This project was funded by a postdoctoral grant to S.D. from the European Union’s Horizon 2020 research and innovation programme under the Marie Skłodowska-Curie grant agreement No 795892. Additional funding came from the Human Frontier Science Program (Career Development Award CDA00047/2017-C to L.-A.C.), the Knut and Alice Wallenberg Foundation (through the Wallenberg Centre for Molecular Medicine Umeå). N.A.-B. and AK were intramurally funded by the National Institutes of Health (USA).

## Author Contributions

S.D., A.K., K.S., N.A.-B., L.-A.C. conceived the project. S.D. performed cryo-electron tomography. A.K. performed virus replication and egress assays. S.D., D.R.M., L.-A.C. performed subtomogram averaging. S.D., A.K., K.S., B.A., D.R.M., N.A.-B., L.-A.C. analyzed data. S.D., L.-A.C. wrote the original draft. All authors reviewed and edited the manuscript.

## METHOD DETAILS

### Cell lines and cultures

HeLa cells were obtained from ATCC (# CRM-CCL-2™). HeLa LC3 and GABARAP 3KO cells were described previously ^35^. All cell lines were grown in (D)MEM supplemented with 10% fetal bovine serum (FBS)/25mM HEPES/GlutaMAX™/ Penicillin-/Streptomycin (Gibco) and maintained at 37ºC in a 5% CO_2_ atmosphere. Cells were regularly screened for the presence of mycoplasma infection.

### Antibodies and virus

Rabbit monoclonal anti-LC3B (D11, Cell Signaling), Mouse monoclonal Anti-β-Actin (A2228, Sigma), Mouse monoclonal anti-VP1(clone B3/H1), Rabbit polyclonal anti-3D gifted by George Belov (UMD), Human anti-A12 gifted by Konstantin Chumakov (FDA). Goat anti-Rabbit IgG (H+L) Cross-Adsorbed Secondary Antibody, Alexa Fluor 555 (A21428, Invitrogen) and Goat anti-Human IgG (H+L) Cross-Adsorbed Secondary Antibody, Alexa Fluor 488 (A11013, Invitrogen). Poliovirus Type 1 Mahoney strain was a gift from George Belov (UMD).

### Drug treatments

HeLa cells were pre-treated with DMSO or 1 μM MRT68921 (SML1644, Merck) in culture medium, 1h before the infection. Medium was replaced with serum free media containing DMSO or 1 μM MRT68921 and cells were infected with Poliovirus (PV) at MOI 5 for 1h. Inoculum was then removed, and cells were incubated in fresh media containing 2% FBS and 1 μM MRT68921 and/or 5μM Vps34-IN1 or 50 μg/mL hydantoin (5-(3,4-dichlorophenyl)-5-methylimidazolidine-2,4-dione (EN300-21815)) for 3h to 8h. Vps34-IN1(17392, Cayman chemicals) was added to the cell media at 1 h p.i. to not interfere with the endocytosis-based viral entry. Hydantoin was added at 1 h p.i. After addition, all inhibitors were kept in the cell media throughout the experiments.

### LC3 lipidation assay

For each condition and time point, HeLa cells were seeded in two T25 flasks (2×10^6^ cells/flask) and infected with poliovirus at MOI 5. Infected cells were collected by trypsinization, then centrifuged at 500 g for 5 min, pellets were resuspended in PBS and washed twice with PBS. Cells were lysed with lysis buffer (20 mM Tris-HCl pH8, 300 mM KCl, 10% glycerol, 0.25% Nonidet P-40, 0.5 mM EDTA, 0.5 mM EGTA) on ice for 15min, passed through a 22G needle and centrifuged at 15,000 rpm for 20 min. After centrifugation, supernatants were stored at - 80°C and protein concentrations were further determined using the Pierce™ BCA Protein Assay Kit (23225, ThermoFisher Scientific).

### SDS-PAGE/Western Blot

Proteins were separated by SDS-PAGE and gels were transferred via a semi-dry blotter to PVDF transfer membranes and blocked for 1 h with TBS-T containing 5% (w/v) milk powder or 5% (w/v) BSA, followed by probing with primary antibodies and overnight incubation, and further re-probed with corresponding HRP conjugated secondary antibodies (Sigma Aldrich) for 1h. Blots were developed using SuperSignal West Pico PLUS Chemiluminescent Substrate (Thermo Scientific) and imaged using the Amersham Imager 600 (GE Healthcare Biosciences) or the Biorad ChemiDoc™ Touch, and analyzed with imageJ.

### Sample preparation for cryo-electron tomography

24h before infection, cells were seeded on R2/2 gold UltrAufoil grids (200 mesh, Quantifoil Micro Tools GmbH, Großlöbichau, Germany) in μ-Slide 8 Well chamber (IBIDI) at 2×10^4^ cells/well. Prior to use, UltrAufoil grids were glow-discharged, dipped in ethanol and then washed with cell media for 30 min. HeLa cells were infected with poliovirus at MOI 5 for 1h in serum-free media at 37°C. The μ-Slide was gently agitated every 15 min to ensure an even coverage and maximize virus contact with the cell monolayer. After 1h of virus absorption, the inoculum was replaced with fresh DMEM media supplemented with 2% FBS. Infected cells were plunge-frozen in liquid ethane-propane mix 3h or 6h post-infection using a Vitrobot plunge freezer (Thermo Fisher Scientific) at 23°C, 90% humidity, with blot force -5 and blot time 6.5 s.

### Cryo-lamella generation and characterization

Cryo-lamellas of poliovirus-infected cells were generated employing the wedge-milling method ^36^ using a Scios focused-ion-beam scanning electron microscope (ThermoFisher Scientific). To prevent sample drift during the milling process and to enhance sample conductivity, the samples were first coated with platinum using the gas injection system (GIS, ThermoFisher Scientific) operated at 26°C, at 12 mm stage working distance and 7 seconds gas injection time. The milling was performed at a tilt angle within a range of 17°– 23° stage tilt. Lamella preparation was performed in a stepwise milling using parallel rectangular pattern above and below the area of interest, with reducing the ion beam current throughout the milling process, from 0.3 nA for the first milling step to remove the bulk material to 0.03 nA at the final cleaning step to obtain the lamella which was set to a minimum thickness of 200 nm. To minimize the contamination of lamellas with ice crystals, they were stored in liquid nitrogen for less than a week before tilt-series collection at the Titan Krios (ThermoFisher Scientific). The final thickness of lamellas was measured at the Titan Krios. Two images of the same area of a lamella were recorded at an intermediate magnification (8700x): an energy-filtered image (F image), and non-filtered image for which the energy filter slit was removed (nF image). The lamella thickness was estimated as 350*ln(I(nF)/I(F)), where I(nF) and I(F) are the intensities in the non-filtered and filtered images, respectively, and 350 the estimate of the electron mean-free path at 300 kV in ice (in nm).

### Cryo-electron tomography

Data was collected using a Titan Krios (ThermoFisher Scientific) operated at 300 kV in parallel illumination mode. Tilt series were recorded using SerialEM software on a K2 Summit detector (Gatan, Pleasanton, CA) operated in super-resolution mode. The K2 Summit detector was mounted on a BioQuantum energy filter (Gatan, Pleasanton, CA) operated with a 20eV slit width. A condenser aperture of 70 μm and an objective aperture of 100 μm were chosen for tilt-series collection. Coma-free alignment was performed with AutoCtf/Sherpa (ThermoFisher Scientific). Tilt-series were acquired in dose-symmetric or bi-directional mode. Due to the pre-tilt of the lamellas, the starting angle used was + 13° for dose-symmetric, and - 21° to - 25° for bi-directional tilt series acquisition. The following parameters were used for acquisition: 33kx nominal magnification corresponding to a specimen pixel size of 2.145 Å; defocus range -3 to -5 μm, tilt-range depending on the lamella pre-tilt and thickness typically ± 50° to ± 60°; tilt increment 2° or 3°; total electron dose ∼100 e−/Å^2^ (bidirectional tilt series) and 130 e-/Å2 (dose-symmetric tilt series). The exposure dose was not varied as a function of tilt angle. At each tilt angle, the exposure was saved as a non-gain-corrected TIFF movie containing a dose per frame of around 0.25 per super-resolution pixel.

### Cryo-electron tomography data processing

Super-resolution TIFF movies were unpacked and gain-reference corrected, and subsequently corrected for sample motion using MotionCor2^37^. The motion correction included a factor 2 binning resulting in a specimen pixel size of 4.29 Å. After reassembly of tilt-series image stacks they were processed using IMOD^38^. Tilt series were aligned using patch tracking. The aligned stacks were CTF-corrected with a custom-made script using CTFFIND4^39^ and CTFPHASEFLIP^40^, a part of IMOD^38^. Tomograms were then generated in IMOD using weighted back projection, no low-pass filtering was performed at this stage. For visualization, tomograms were 4 times binned using IMOD, resulting in a pixel size of 17.16 Å and denoised using the boxfilter option in Amira software (ThermoFisher Scientific). Filtered tomograms were further segmented using Amira for the representation of membranes, protein densities and filaments, the last using the Fiber tracing module. Subtomogram averages of empty capsids and RNA-loaded virions were integrated in these 3D renderings through UCSF Chimera^41^.

### Subtomogram averaging of viruses

Subtomogram averaging of virus particles for visualization purposes was performed using Dynamo ^42,43^. For RNA-loaded virions, a total of 300 particles were manually picked in two tomograms containing the cytoplasmic virus arrays (MRT68921-treated cells). For empty viruses, a total 198 particles were manually picked in two tomograms from Hydantoin-treated cells. Subvolumes of unbinned particles (4.29 Å/px) were extracted with a box size of 100*100*100 voxels. Subvolumes were first iteratively aligned against each other allowing only shifts using a spherical mask with radius of 48px and a Gaussian falloff of 3 px. After centering particles, a full rotational alignment was performed. The average was realigned to icosahedral symmetry axes and a final average was calculated with imposed icosahedral symmetry. Gold-standard FSC curves for resolution estimation of the virus averages were calculated in Dynamo, resulting in a resolution estimate of 25 Å for empty capsids and 30 Å for RNA-loaded virions at a Fourier shell correlation threshold of 0.143. Subtomogram averages were low-pass filtered to their respective resolution and the 3D renderings were created in UCSF Chimera.

### Subtomogram averaging of filaments

The subtomogram selection, extraction and averaging followed the workflow schematically presented in Fig. S8. 1856 filaments were traced in 16 tomograms using the fiber tracing module ^44^ in Amira (Thermo Fisher Scientific). The tracing was performed within manually segmented regions corresponding to filament-filled autophagy-like membranes, and the tracing parameters were selected so that the number and length of the filaments was consistent with their visual appearance in the tomograms. Filament coordinates were exported from Amira and imported to Dynamo using a custom-written MATLAB (Mathworks) script, after which Dynamo was used to extract subtomograms at regular intervals. Initially, an oversampled subtomogram extraction was performed, with (100px)^3 subtomograms extracted along the filament axis at 5 px intervals. This resulted in 55,376 subtomograms. Each subtomogram was assigned initial Euler angles that aligned it with the traced filament axis, but gave it a randomized rotation along that axis. The subsequent alignments and classifications were performed using a cylindrical mask with a radius of 16 px and a Gaussian falloff of 3 or 5 px. Subtomogram averaging was performed in Dynamo. After an initial single iteration allowing for centering of the subtomograms perpendicularly to the filament axis, several iterations were performed allowing for shifts, free rotations around the filament axis and a +/-30° tilt with respect to the filament axis. At this stage of the data processing, efforts to determine filament polarity by allowing each subtomogram in a given filament to change orientation during alignment, and then imposing the majority orientation on all subunits of the filament were not yet successful. From the full oversampled dataset, overlapping subtomograms were removed using a distance threshold of 10 px (42.9 Å), resulting in a set of 16,682 subtomograms. These subtomograms were subjected to a five-class multireference alignment (MRA) in Dynamo.

Three of the five classes had a similar, regular helical appearance, and were pooled for a second round of MRA with five classes, from which one class of 3517 subtomograms was better defined than the others. This class was subjected to a half-set gold-standard refinement in Dynamo resulting in a resolution estimate of 23.8 Å at a Fourier shell correlation threshold of 0.143. The resulting average allowed a first estimation of the helical parameters as a rise of ∼56 Å per subunit and a rotation of ∼36° per subunit. At this point, tomograms were re-reconstructed using NovaCTF to allow for 3D CTF correction by phase flipping^45^. Subtomograms were reextracted at positions corresponding to the helical subunits, which after removal of overlapping particles resulted in an enlarged data set of 10897 subtomograms. These subtomogram poses and positions were exported from Dynamo for further processing in the subTOM package written in MATLAB with functions adapted from the TOM^46^, AV3^47^ and Dynamo packages. The scripts and relevant documentation are available to download [https://www2.mrc-lmb.cam.ac.uk/groups/briggs/resources]. Additionally, instead of a binary wedge mask, a modified wedge mask was used^48^. The missing wedge was modelled at all processing stages as the average of the amplitude spectra of subtomograms extracted from regions of each tomogram containing empty ice, and was applied during alignment and averaging. We applied both principal component analysis (PCA) and multivariate statistical analysis (MSA) to classify subtomograms by filament straightness and similar helical parameters. The PCA was performed on wedge-masked difference (WMD) maps^49^ with calculations implemented in MATLAB using code adapted from PEET^49^ and Dynamo packages, and MSA also implemented in MATLAB using code adapted from I3^50^. Movies detailing the variability related to each Eigenvolume from MSA classification were generated using Eigen-filtering / reconstruction methods, and similar movies from WMD classifications were generated by producing class averages sorted by determined Eigencoefficients. Improved class averages allowed for recalculation of helical parameters and further sub-boxing as well as determining the filament polarity. Specifically, the sub-box poses and positions were determined by auto-correlation of the reference after rotation along the filament axis, which yielded a helical symmetry with a rise of 51.5 Å and rotation of 29°, and then duplicates within a sphere of diameter 50 Å were removed. Poses were not adjusted from their gold-standard alignment values and the Fourier shell correlation determined resolution was improved to 18.5 Å at the 0.143 threshold. Following the FSC calculation, the half-maps were averaged, filtered to the measured resolution by the determined FSC-curve and sharpened using a heuristically determined B-factor of -4000 Å^2 51^.

### Quantitative analysis of membrane structures and virions

Tomograms were visually inspected and membrane structures such as single-membrane vesicles and tubes were assigned as single membranes. Double membrane structures were assigned as phagophore-like membranes if they had a clear opening within the tomogram volume (i.e. were cup-shaped), and as double-membrane vesicles if they did not have an opening. Collectively, these two types of double-membrane structures were referred to as autophagy-like membranes. Virus particles were localized and counted in each tomogram with template matching using PyTom ^52^. For empty capsids and RNA-loaded virions template matching was performed using respectively the re-sampled, filtered empty capsid (EMD-9644) and mature virion (EMD-9642) cryo-EM structures of the human coxsackievirus A10 ^53^. The concentration of each structures was obtained by dividing the total number of structures by the volume of the corresponding tomogram. The number of tomograms obtained for all the conditions are listed in the supplementary table S1. The closure of viral capsid intermediates was calculated using IMOD. First, in a central section through the capsid intermediate, the circumference was traced and measured with IMOD drawing tools, then the open length was divided by the circumference of a complete virus particle. This fraction was multiplied by 360 to get the angle of the cone that describes the capsid assembly intermediate.

### Freeze substitution

Poliovirus-infected HeLa cells non-treated and treated with MRT68921 were grown on carbon-coated sapphire discs and high-pressure frozen at 6 h p.i. using a Leica HPM100. Freeze substitution was performed as a variation of the Kukulski protocol ^54^. Briefly, sapphire discs placed in adapted carriers were filled with freeze substituent (0.1% uranyl acetate in acetone, 1% H2O) and placed in a temperature-controlling AFS2 (Leica) equipped with an FPS robot. In the first step, freeze-substitution occurred at -90ºC for 48 h then the temperature was raised to -45ºC. The samples were maintained in the freeze substituent at -45ºC for 5 h before washing 3 times with acetone followed by a temperature increase and infiltration with increasing concentrations of Lowicryl HM20. Finally, the samples were gradually warmed up to -25°C before infiltrating 3 times with 100% Lowicryl and UV-polymerized for 48h at -25ºC. Polymerization then continued for another 24 h at room temperature. The embedded samples were sectioned with a diamond knife (DiATOME-90) to slices of 60-120 nm thickness by using ultramicrotome (Reichart ULTRACUT S). Imaging was performed in FEI Talos electron microscope operating at 120 kV. Grids were examined at a Talos L120C (FEI, Eindhoven, The Netherlands) operating at 120kV. Micrographs were acquired with a Ceta 16M CCD camera (FEI, Eindhoven, The Netherlands) using TEM Image & Analysis software ver. 4.17 (FEI, Eindhoven, The Netherlands).

### Confocal microscopy analysis

HeLa wildtype (WT), LC3 3KO and GAB 3KO cells were infected with PV for 1h at MOI 5 in serum free media, washed and kept in 2% FCS DMEM/high glucose for 3h or 6h. Cells were fixed in 4% paraformaldehyde (PFA)/phosphate buffer solution (PBS) for 10 min at room temperature. Primary and secondary antibody incubations were carried out in PBS/10%FBS supplemented with saponin at 0.2% for 1h at room temperature. Cells were rinsed twice in PBS, twice in water and mounted with Dapi Fluoromount-G (Electron Microscopy Science). Image acquisition was performed using a LSM780 confocal microscope (Carl Zeiss) with a 63X/1.4 NA oil objective or a 40X/1.4 NA oil objective. Cell numbers and individual cells respective mean fluorescence intensities and area were obtained using ImageJ.

### Virus titration analysis of cell supernatants

Extracellular medium was collected from infected cell cultures and serially diluted in 96 well plates (10^−1^ to 10^−8^). Dilutions were subsequently used to inoculate in triplicates HeLa WT cells seeded at 6×10^4^ cells/well in 96-well plates. Cells were incubated at 37°C for 44h, fixed with 10% paraformaldehyde and stained with crystal violet. TCID50/ml was calculated using the Spearman & Kärber algorithm.

### Vesicle isolation

HeLa wildtype (WT), LC3 3KO and GAB 3KO cells were infected with PV for 1h at MOI 5, washed and kept in serum-free DMEM/high glucose for 8h. Supernatants obtained from one well of a 6-well plate (1.5mL, ∼500000 cells) were harvested and centrifuged at 1000g for 10min, then at 20000g for 30min. Pellets were either resuspended in 1X loading buffer and analyzed by Western Blot, or resuspend in RNA lysis buffer and processed for RT-PCR.

### Quantitative (q) PCR analysis

Cell supernatants and cell lysates were harvested at specific time points and lysed using RNA lysis buffer provided in the RNA isolation kit (Quick-RNA Microprep Kit, Zymo Research). RNA isolation was performed as per the manufacturer’s instructions and cDNA was prepared using Thermo Scientific Maxima First Strand cDNA Synthesis Kit for RT-qPCR (Fisher Scientific). RT-PCR was performed using iTaq Universal SYBR® Green Supermix (BioRad) in Roche LightCycler 96 system (Roche), using the following thermal cycling conditions: 95°C for 90s, 40 cycles at 95°C for 10 sec, 57°C for 10 sec and 72°C for 110 sec. The samples were run in duplicate for each data point. Primers used: For 5 ‘CGGCTAATCCCAACCTCG 3’, Rev 5’ CACCATAAGCAGCCACAATAAAATAA 3’.

### Quantification and statistical analysis

Data and statistical analysis were performed using Prism (GraphPad Software Inc., USA). Details about replicates, statistical test used, exact values of n, what n represents, and dispersion and precision measures used can be found in figures and corresponding figure legends. Values of p < 0.05 were considered significant.

## Data and code availability

Cryo-EM map of the filament structure has been deposited in the Electron Microscopy Data Bank (EMDB) under accession code EMD-XXXX. Any additional information required to reanalyze the data reported in this paper is available from the lead contact upon request.

**Table S1:**
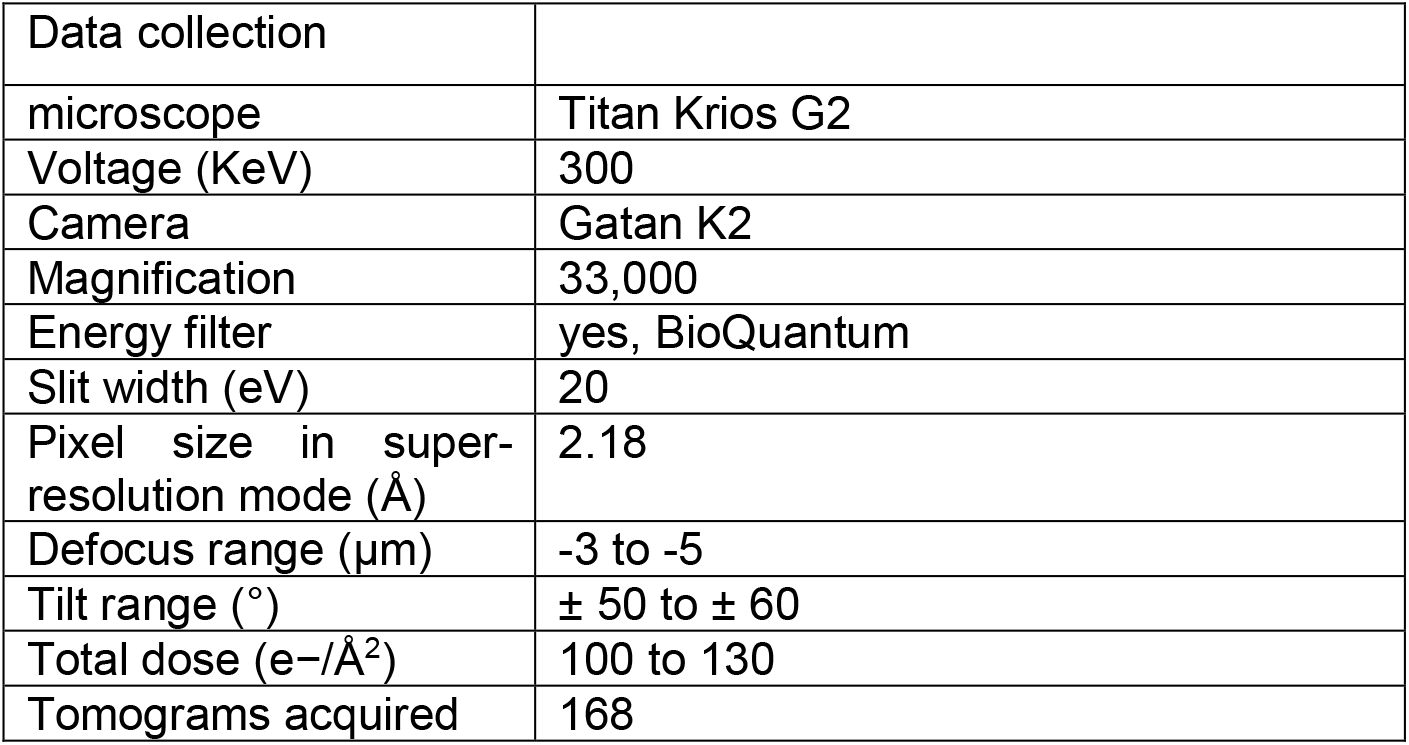
Data collection and parameter for Cryo-electron tomography of poliovirus replication and assembly sites.

**Table S2:**
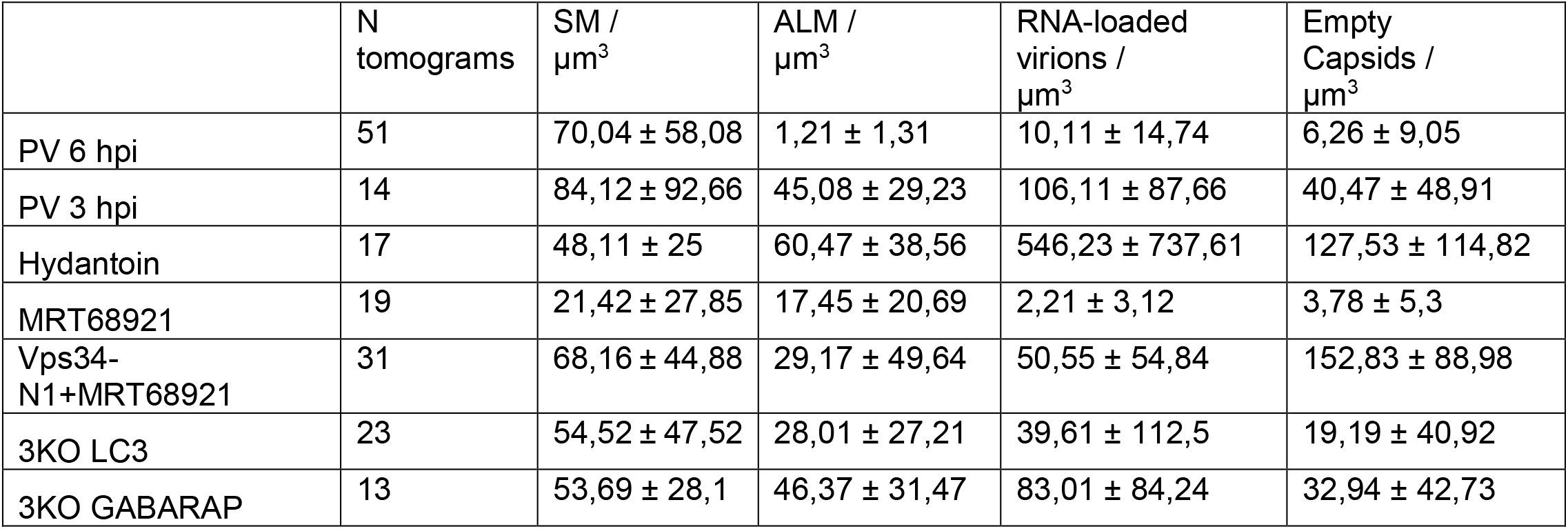
Concentration of membrane structures and virions (Mean ± SD).

**Table S3:**
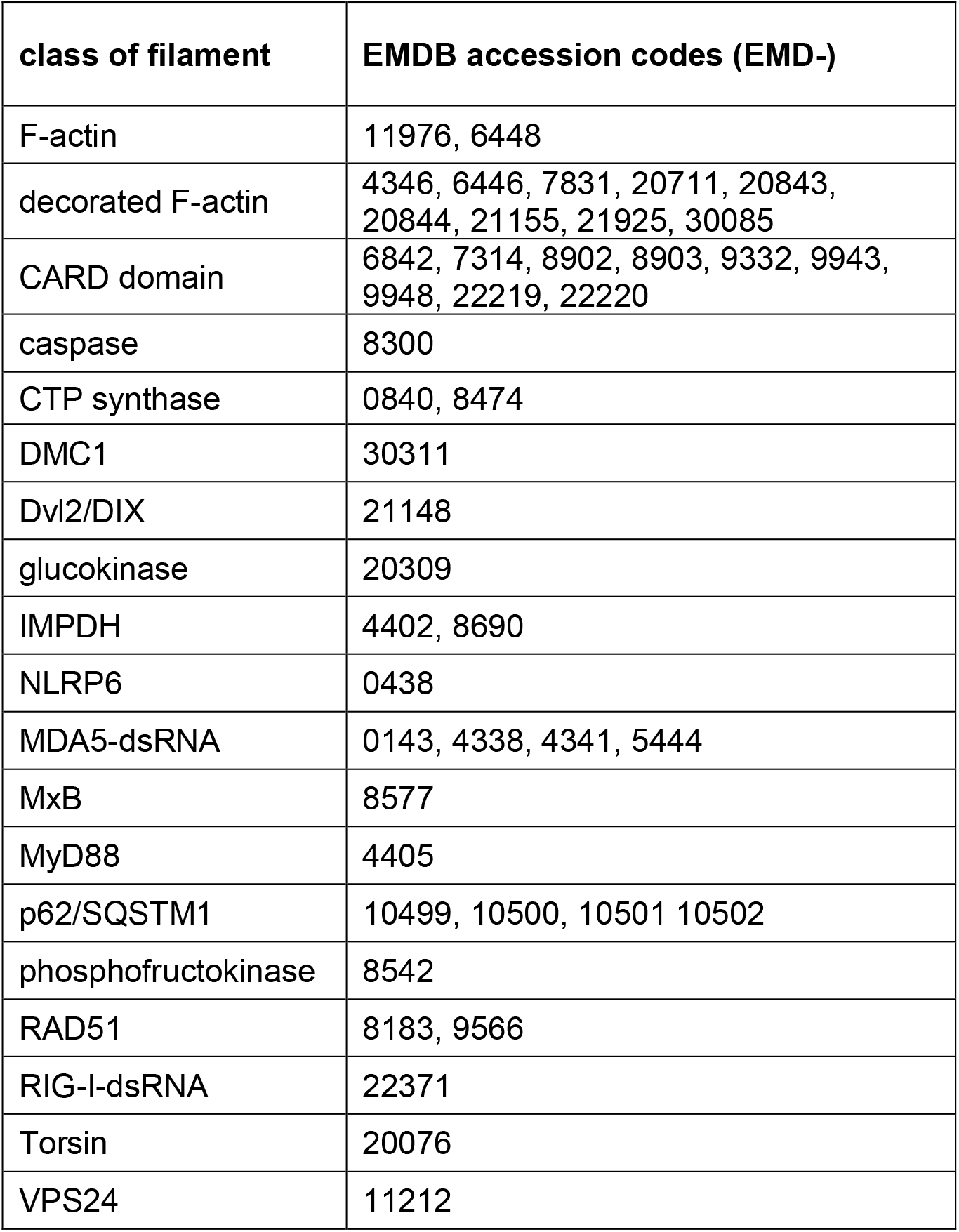
Protein filament structures from the electron microscopy data base compared to the ALM-associated filament. The filaments are listed by type. Fig. S7 shows the subset of filaments being most relevant either by biological function or close structural match.

---

**Figure S1:**
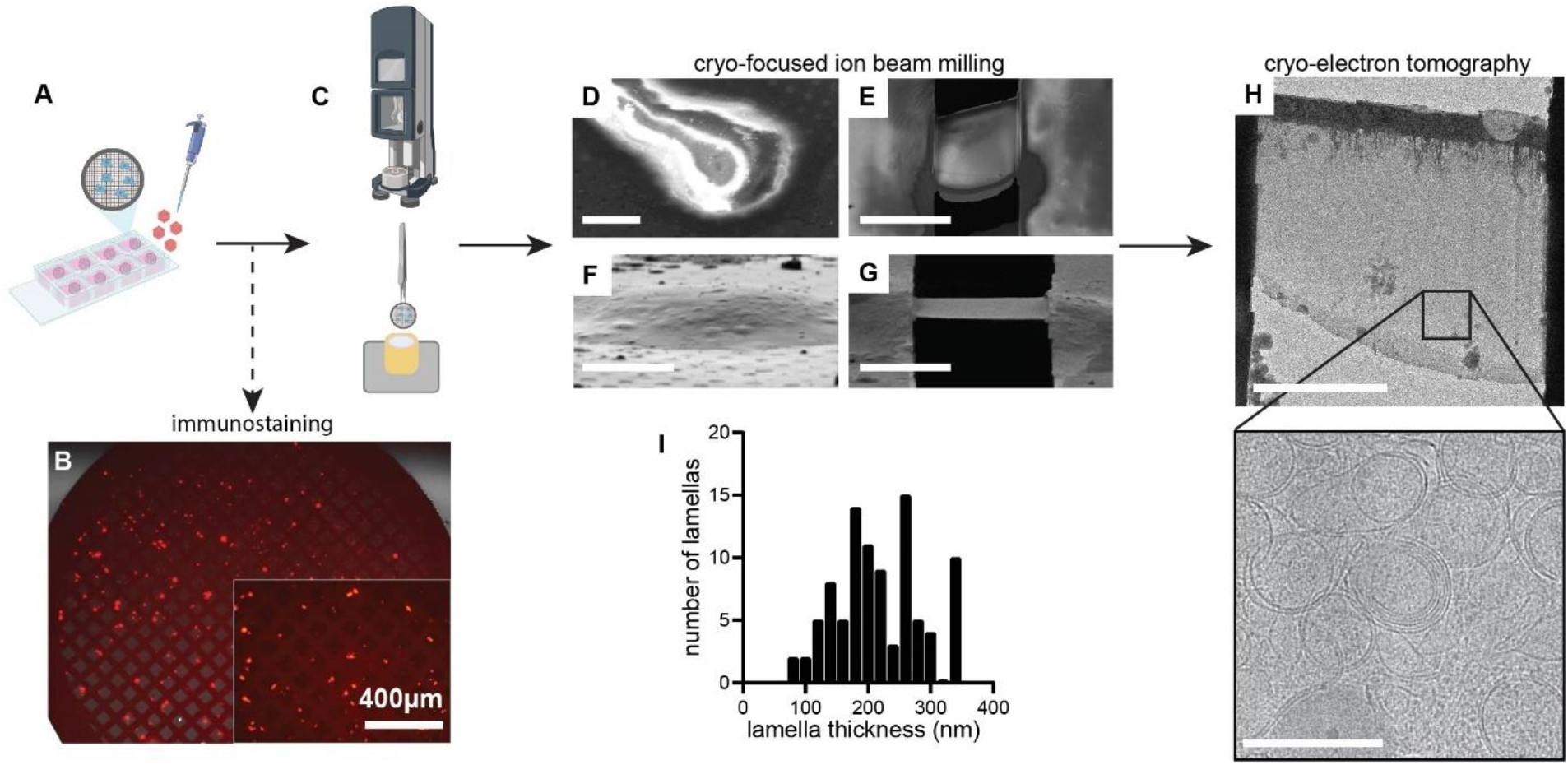
Experimental workflow for cryo-electron tomography. (A) HeLa cells were cultured on gold EM grids in chambered slides and infected with poliovirus at MOI 5. (B) Immunostaining against the viral capsid protein VP1, with a secondary antibody conjugated to Alexa584 (red), confirmed PV infection of virtually all cells growing on EM grids. (C) Unfixed and unlabeled infected cells were vitrified by plunge freezing. (D-E) Scanning electron microscopy and (F-G) ion beam images before and after the FIB-milling process of a PV-infected cell, resulting in thin lamella (here seen at a slight angle). (H) Low magnification transmission electron microscopy (TEM) image of the thin transparent cryo-lamella, where areas containing DMVs were abundant (zoomed box). (I) Thickness distribution of cryo-lamellas obtained with TEM. Average thickness is 217 ± 69 nm; n = 93 lamellas. Scale bars: (F-G) 5 μm, (H) 200 nm.

**Figure S2:**
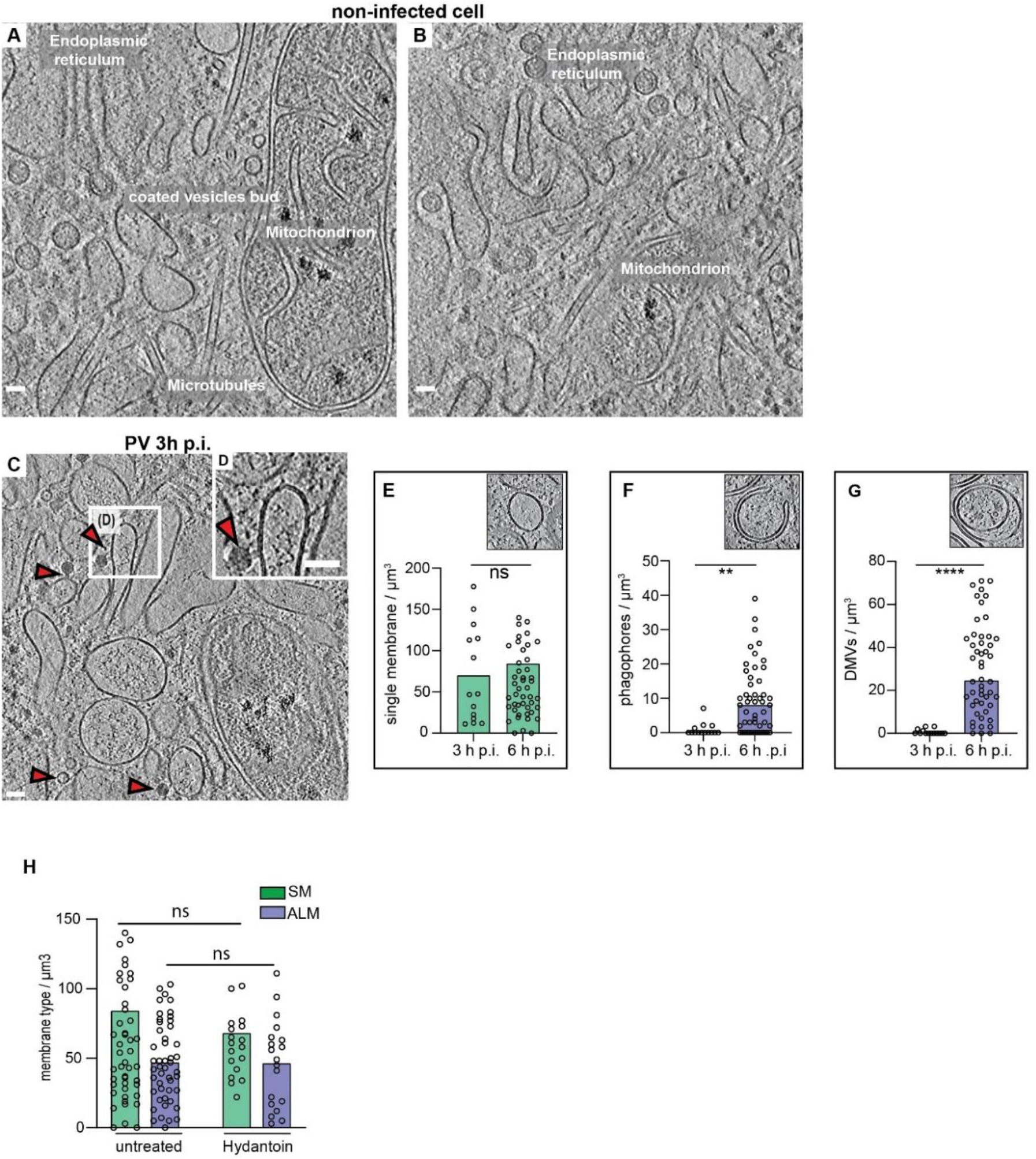
Cryo-ET of poliovirus-infected cell at 3 h p.i. (A-B) Slices through a tomogram of an uninfected HeLa cell showing several cytoplasmic features and organelles as labeled. (C) Slice through a cryo-electron tomogram of a lamella milled through a PV-infected cell at 3 h p.i., revealing PV-induced SM proliferation. (D) Magnified view of the white box in (C) showing a tethered virion to SM. (C-D) Red arrowheads indicate viral particles. Scale bars: 50 nm. (E-G) Scatter plots with bars representing the mean concentration of SM structures (E) phagophore-like structures (F), and DMVs (G), observed in cryo-tomograms of 3 h p.i. and 6 h p.i. Insets: slices through different tomograms showing the type of membrane for each graph. (H) Scatter plot with bars representing the mean concentration of SM and ALM measured in Hydantoin-treated cells at 6 h p.i. compared to untreated cells. Each dot corresponds to one tomogram, bars represent the averages (see also Supplementary table 2).

**Figure S3:**
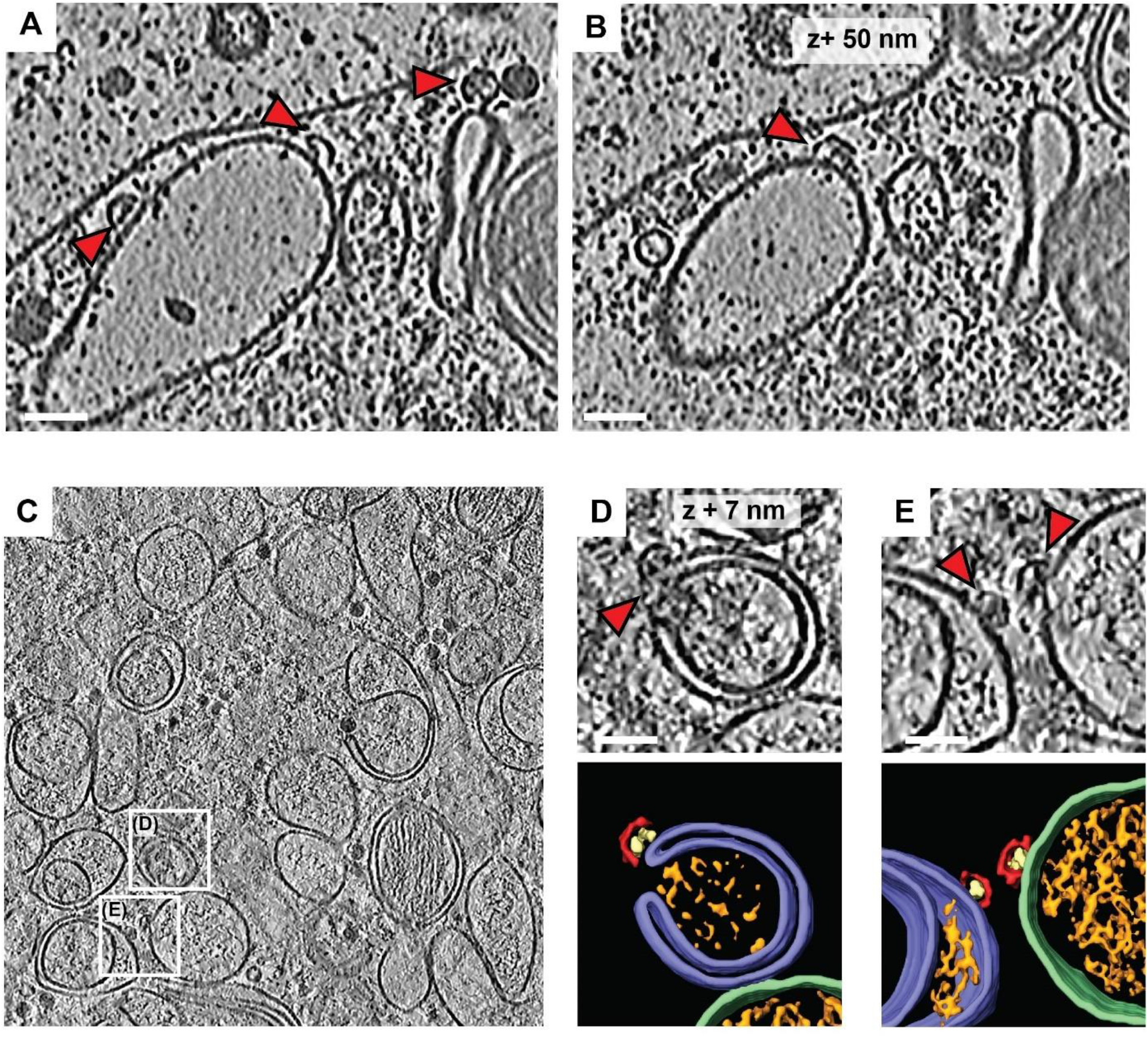
capsid intermediates bound to SM and ALM. (A-B) Slices of PV-infected cell at 6 h p.i. cryo-tomogram corresponding to the segmentation in Figure 2A, showing capsid intermediates bound to SM (red arrowheads). (C) Cryo-electron tomogram of a PV-infected cell at 6 h p.i. White boxes indicate areas with capsid assembly intermediates. (D-E) Magnified views of boxes in (C) with capsid intermediates indicated by red arrowheads, and their corresponding 3D segmentations showing capsid assembly intermediates (red) containing luminal densities (yellow), associated with autophagy-like membranes (ALMs, purple) and a single-membrane vesicle (SM, green). Scale bars 50 nm.

**Figure S4:**
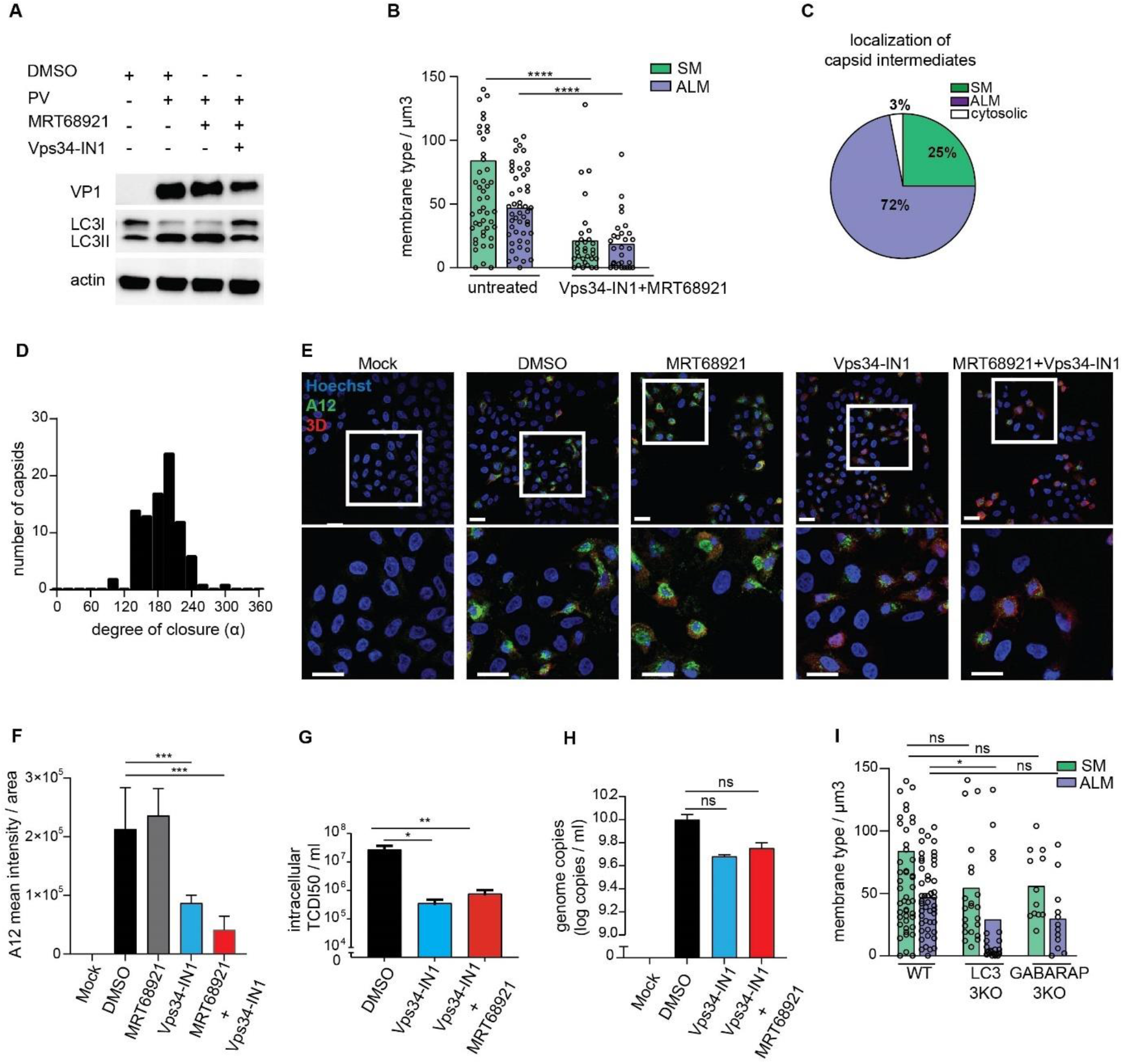
Characterization of PV-infection in Vps34-IN1 treated cells. (A) LC3B lipidation in PV-infected cells at 6 h p.i. treated or not with the autophagy inhibitors MRT68921 (1 μM) and/or Vps34-IN1 (5 μM). Cell lysates were immunoblotted against the indicated proteins. LC3II, lipidated form of LC3B protein. (B) Concentration of SMs and ALMs measured in DMSO and Vps34-IN1 + MRT68921 treated cells at 6 h p.i. (C) Percentage of capsid assembly intermediates found on SMs, ALMs or not associated with membranes, as counted in 17 cryo-tomograms of Vps34-IN1 treated cells at 6 h p.i. (D) Distribution of capsid intermediate closures observed in tomograms of Vps34-IN1 + MRT68921-treated cells at 6 h p.i. The average closure is 186° (SD=35°, N=90). (E) Immunofluorescence assay of PV-infected cells in the presence or absence of autophagy inhibitors MRT68921 and Vps34-IN1 at 6 h p.i. Assembled provirions and mature viruses were detected using the A12 antibody (green) and the expression of the non-structural protein 3D^pol^ was verified concomitantly (red). (F) Quantification of the A12 mean fluorescence intensities obtained from immunofluorescence images illustrated in (B). Bars represent the means ± SD. (G) Intracellular PV titers measured at 6 h p.i. in the presence of Vps34-IN1 and MRT68921. Bars represent the means of biological triplicates ± SEM. (H) Intracellular viral RNA copies measured at 6 h p.i. in cells treated with DMSO, Vps34-IN1 and Vps34-IN1 + MRT68921. (I) Concentration of SMs and ALMs measured in WT, LC3 and GABARAP 3KO cells at 6 h p.i. (B-I) Each dot is one tomogram, the bars represent the average. Numerical source data are presented in Supplementary Table 2. (-D-G) Bars represent the means of biological triplicates ± SEM. Statistical significance by unpaired two-tailed Student’s t test; *p<0,05; **p<0,01 and ***p <0,001. Scale bars: 20 μm.

**Figure S5:**
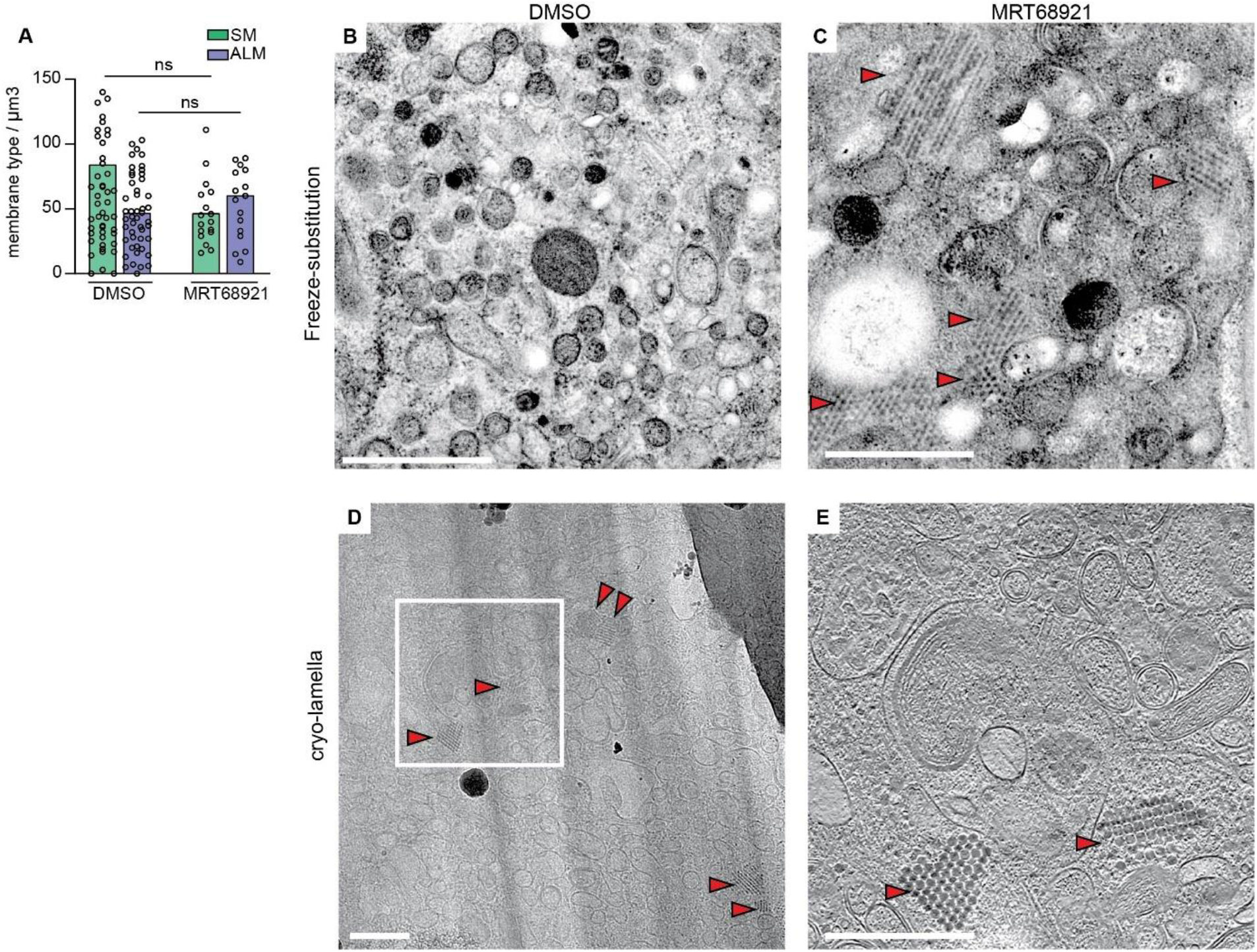
Intracellular virus array formation in infected cells treated with MRT68921. (A)Concentration of SMs vs. ALMs observed in cryo-tomograms of MRT68921 treated cells compared to untreated cells at 6 h p.i. Each dot corresponds to one tomogram analyzed and bars represent the mean (see also Supplementary table 2). (B-C) Electron micrographs of thin sections of freeze-substituted PV-infected cells in the absence (B) and presence (C) of MRT68921 at 6 h p.i., indicating the formation of virus arrays (red arrowheads) when cells are treated with MRT68921. (D) Low magnification image of cryo-lamella milled through MRT68921 treated cell at 6 h p.i. is included for comparison with (C), in which virus arrays are already identified (red arrowhead). (E) Corresponding slice through the tomogram reconstructed from tilt-series collected on the indicated region (white frames in (C)). Scale bars: 500 nm.

**Figure S6:**
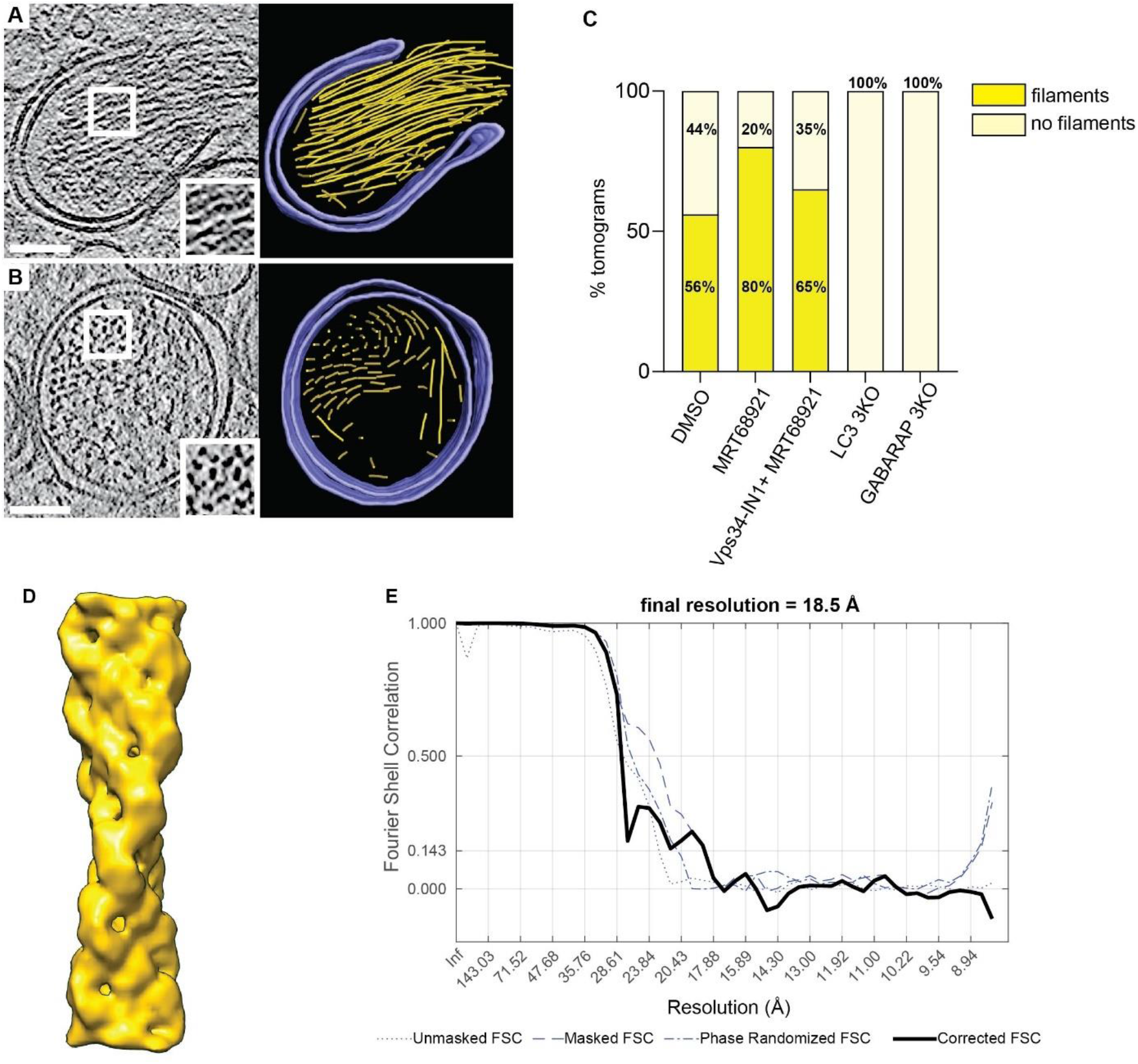
ALM-associated protein filament. (A-B) Slices through tomograms of PV-infected cells at 6 h p.i and their corresponding 3D segmentation showing large bundles of the filament (yellow) filling the interior of a phagophore-like structure (A) and double-membrane vesicle (B) (purple). Zoomed regions (white boxes) highlight the helical twist (A) and bundle formation (B) of the filaments. Scale bars: 100 nm. (C) Percentage of tomograms containing ALM-associated protein filament bundles in DMSO treated WT cells (N=34), in MRT68921 treated WT cells (N=10) and in MRT68921 + Vps34-IN1 treated WT cells (N=20), compared to LC3 (N=17) and GABARAP (N=13) 3KO cells. (D) Subtomogram average of the filaments at 18.5 Å resolution. (E) Fourier shell correlation curves for unmasked, masked, and phase-randomized (beyond 31 Å) half-sets. The corrected curve, equalling 0.143 at 18.5 Å resolution, is based on Chen *et al*^55^ as implemented in subTOM.

**Figure S7:**
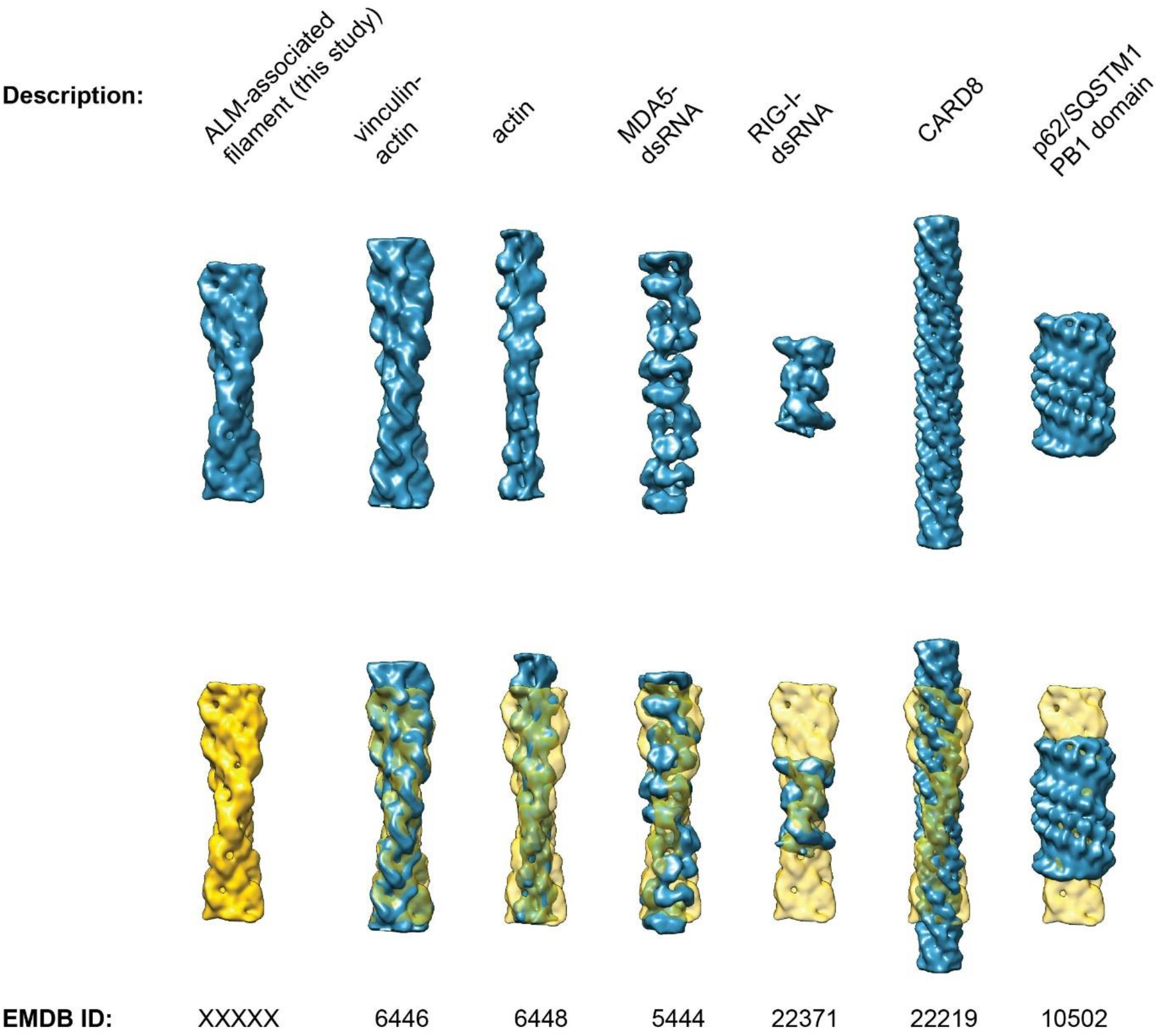
Comparison of the ALM-associated filament with a selection of known filament structures. Top row: isosurface representations of the subtomogram average of the ALM-associated protein filament from this study, and one representative of each of several classes of cellular protein filaments with known structure. In the lower row, the ALM-associated filament is shown in yellow, and semitransparent yellow when fitted to each of the other filaments using UCSF Chimera’s Fit in Map function^41^. Prior to comparison, all filaments were resampled to the same voxel size and filtered to 19 Å resolution. EMDB identifiers are indicated below each volume.

**Figure S8:**
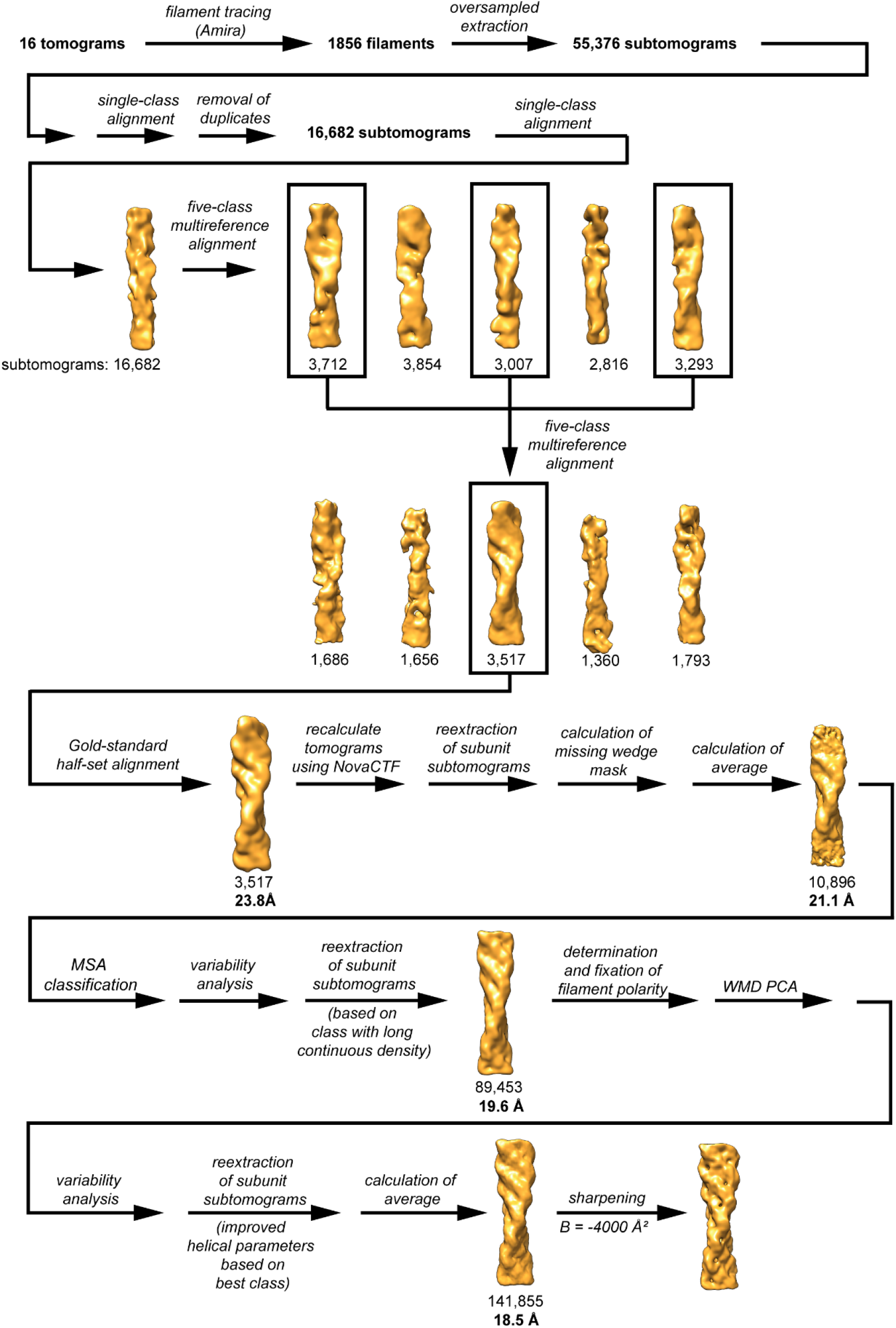
Schematic of the workflow for subtomogram averaging of the ALM-associated filament. The schematic illustrates the major steps of the data processing starting at filament tracing in the tomograms, and extraction of subtomograms along the filament axis, through the several steps of classification, reextraction and averaging.

