## Supplementary Information for "Membrane-assisted assembly and selective autophagy of enteroviruses"

**Supplemental Information**

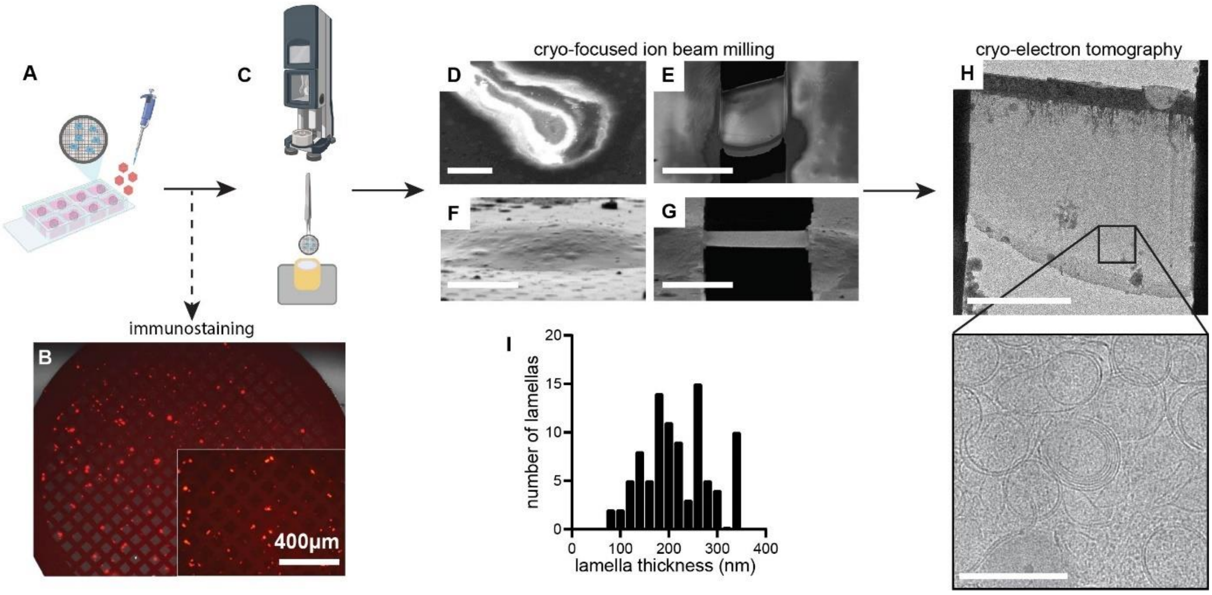


**Figure S1: Experimental workflow for cryo-electron tomography.** (A) HeLa cells were cultured on gold EM grids in chambered slides and infected with poliovirus at MOI 5. (B) Immunostaining against the viral capsid protein VP1, with a secondary antibody conjugated to Alexa584 (red), confirmed PV infection of virtually all cells growing on EM grids. (C) Unfixed and unlabeled infected cells were vitrified by plunge freezing. (D-E) Scanning electron microscopy and (F-G) ion beam images before and after the FIB-milling process of a PV-infected cell, resulting in thin lamella (here seen at a slight angle). (H) Low magnification transmission electron microscopy (TEM) image of the thin transparent cryo-lamella, where areas containing DMVs were abundant (zoomed box). (I) Thickness distribution of cryo-lamellas obtained with TEM. Average thickness is 217 ± 69 nm; n = 93 lamellas. Scale bars: (F-G) 5 µm, (H) 200 nm.

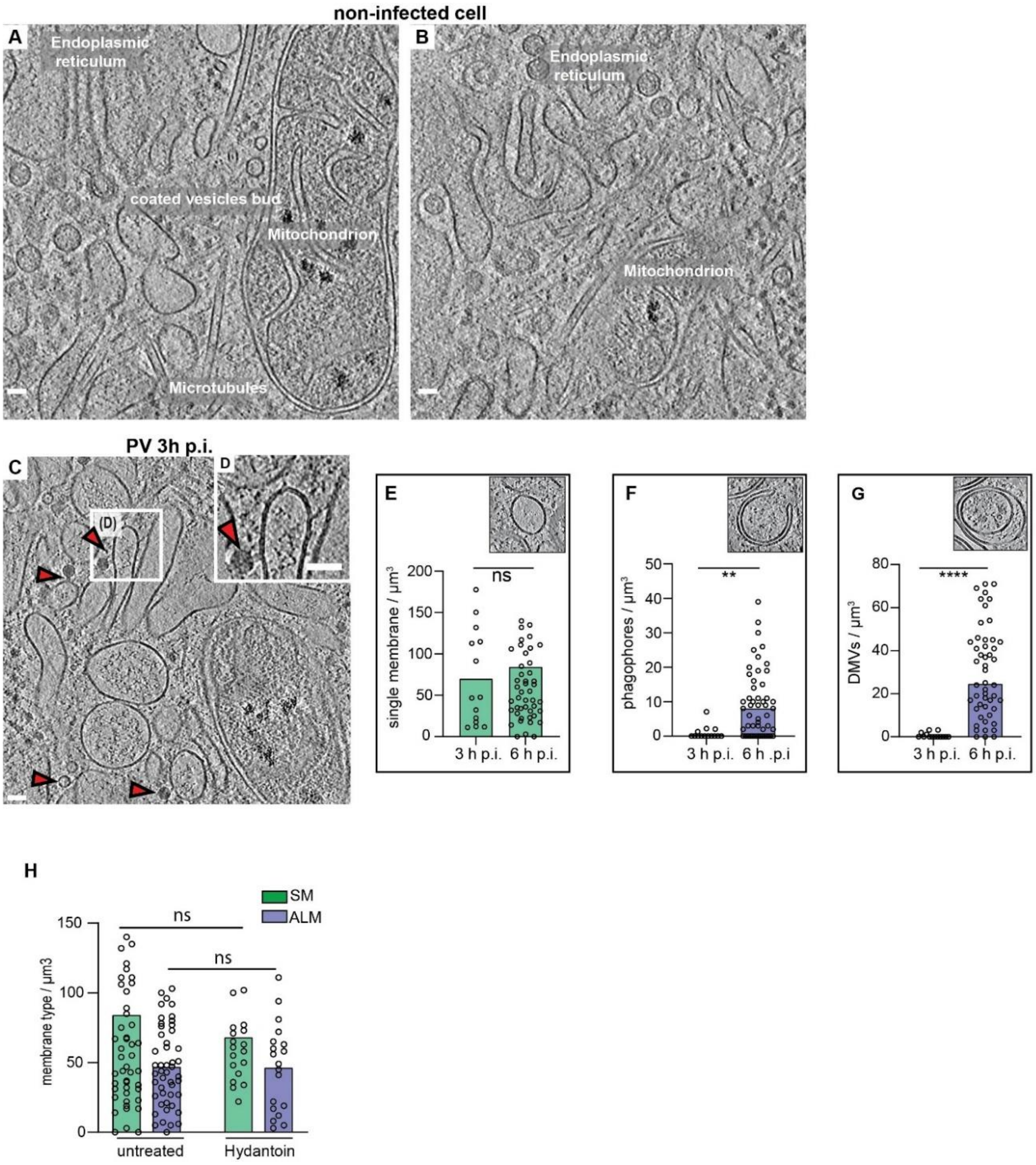


**Figure S2: Cryo-ET of poliovirus-infected cell at 3 h p.i.** (A-B) Slices through a tomogram of an uninfected HeLa cell showing several cytoplasmic features and organelles as labeled. (C) Slice through a cryo-electron tomogram of a lamella milled through a PV-infected cell at 3 h p.i., revealing PV-induced SM proliferation. (D) Magnified view of the white box in (C) showing a tethered virion to SM. (C-D) Red arrowheads indicate viral particles. Scale bars: 50 nm. (E-G) Scatter plots with bars representing the mean concentration of SM structures (E) phagophore-like structures (F), and DMVs (G), observed in cryo-tomograms of 3 h p.i. and 6 h p.i. Insets: slices through different tomograms showing the type of membrane for each graph. (H) Scatter plot with bars representing the mean concentration of SM and ALM measured in Hydantoin-treated cells at 6 h p.i. compared to untreated cells. Each dot corresponds to one tomogram, bars represent the averages (see also Supplementary table 2).

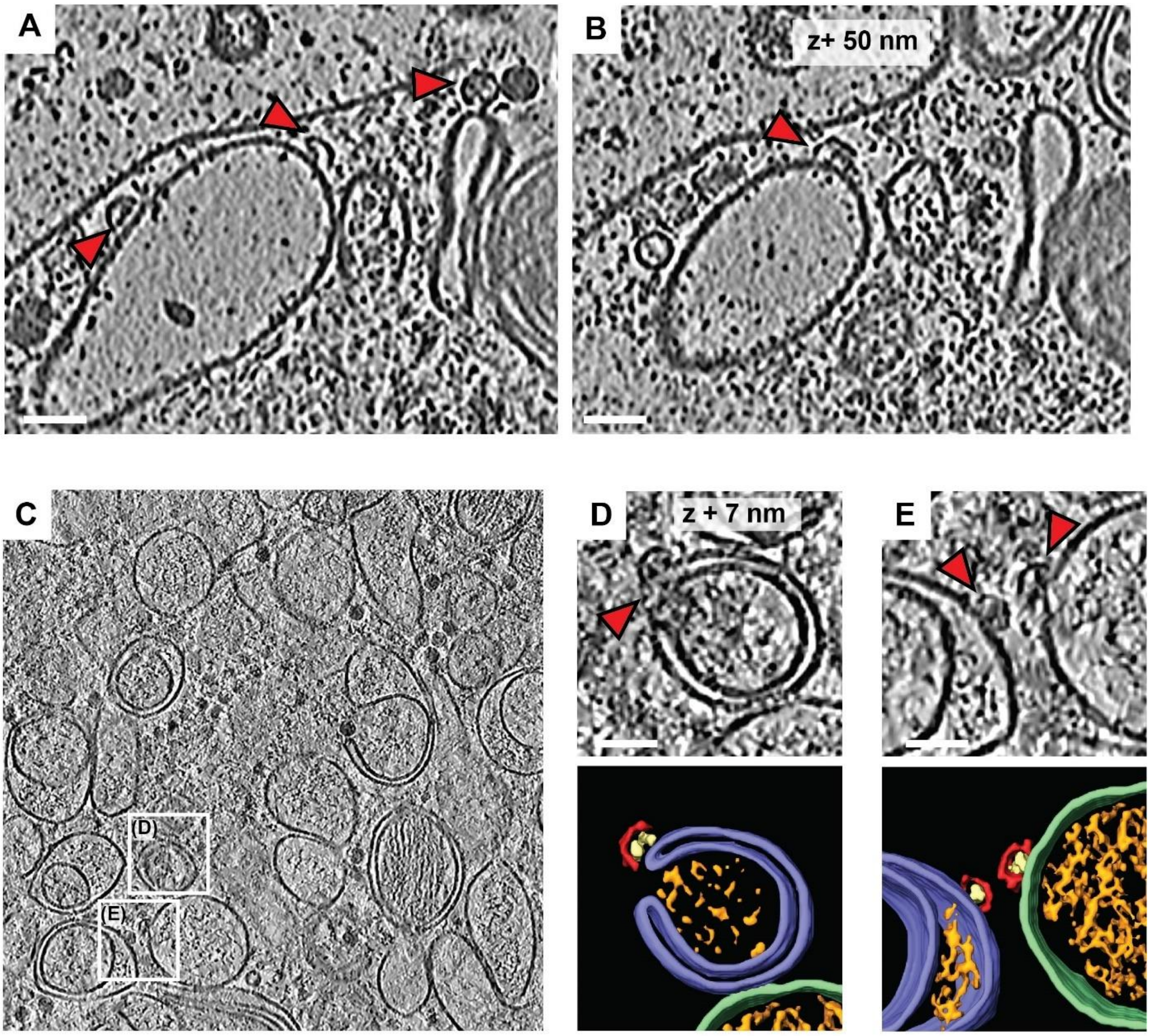


**Figure S3: capsid intermediates bound to SM and ALM**. (A-B) Slices of PV-infected cell at 6 h p.i. cryo-tomogram corresponding to the segmentation in Figure 2A, showing capsid intermediates bound to SM (red arrowheads). (C) Cryo-electron tomogram of a PV-infected cell at 6 h p.i. White boxes indicate areas with capsid assembly intermediates. (D-E) Magnified views of boxes in (C) with capsid intermediates indicated by red arrowheads, and their corresponding 3D segmentations showing capsid assembly intermediates (red) containing luminal densities (yellow), associated with autophagy-like membranes (ALMs, purple) and a single-membrane vesicle (SM, green). Scale bars 50 nm.

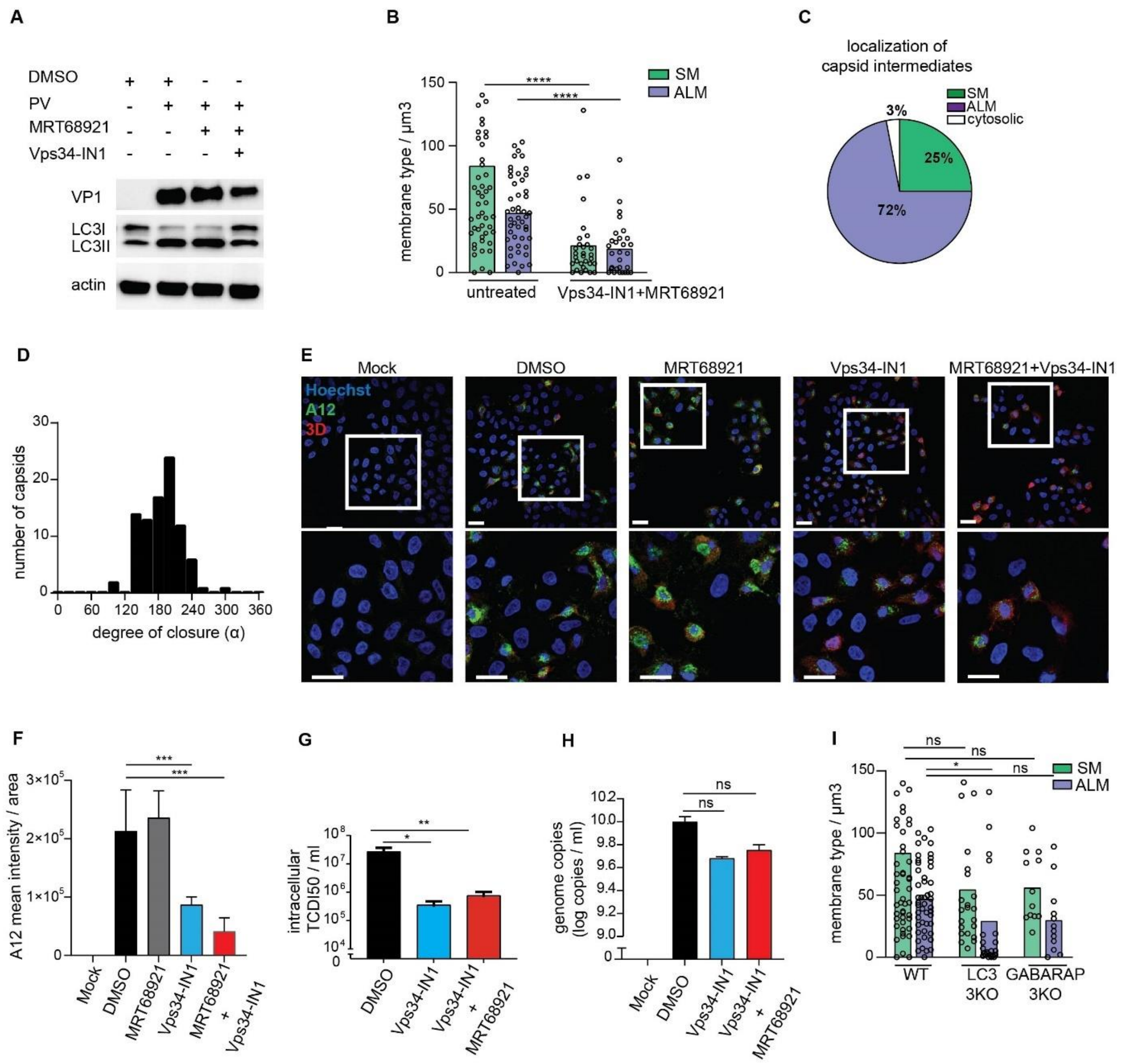


**Figure S4: Characterization of PV-infection in Vps34-IN1 treated cells.** (A) LC3B lipidation in PV-infected cells at 6 h p.i. treated or not with the autophagy inhibitors MRT68921 (1 µM) and/or Vps34-IN1 (5 µM). Cell lysates were immunoblotted against the indicated proteins. LC3II, lipidated form of LC3B protein. (B) Concentration of SMs and ALMs measured in DMSO and Vps34-IN1 + MRT68921 treated cells at 6 h p.i. (C) Percentage of capsid assembly intermediates found on SMs, ALMs or not associated with membranes, as counted in 17 cryo-tomograms of Vps34-IN1 treated cells at 6 h p.i. (D) Distribution of capsid intermediate closures observed in tomograms of Vps34-IN1 + MRT68921-treated cells at 6 h p.i. The average closure is 186° (SD=35°, N=90). (E) Immunofluorescence assay of PV-infected cells in the presence or absence of autophagy inhibitors MRT68921 and Vps34-IN1 at 6 h p.i. Assembled provirions and mature viruses were detected using the A12 antibody (green) and the expression of the non-structural protein $3D^{pol}$ was verified concomitantly (red). (F) Quantification of the A12 mean fluorescence intensities obtained from immunofluorescence images illustrated in (B). Bars represent the means ± SD. (G) Intracellular PV titers measured at 6 h p.i. in the presence of Vps34-IN1 and MRT68921. Bars represent the means of biological triplicates ± SEM. (H) Intracellular viral RNA copies measured at 6 h p.i. in cells treated with DMSO, Vps34-IN1 and Vps34-IN1 + MRT68921. (I) Concentration of SMs and ALMs measured in WT, LC3 and GABARAP 3KO cells at 6 h p.i. (B-I) Each dot is one tomogram, the bars represent the average. Numerical source data are presented in Supplementary Table 2. (-D-G) Bars represent the means of biological triplicates ± SEM. Statistical significance by unpaired two-tailed Student's t test; *p<0,05; **p<0,01 and ***p <0,001. Scale bars: 20 µm.

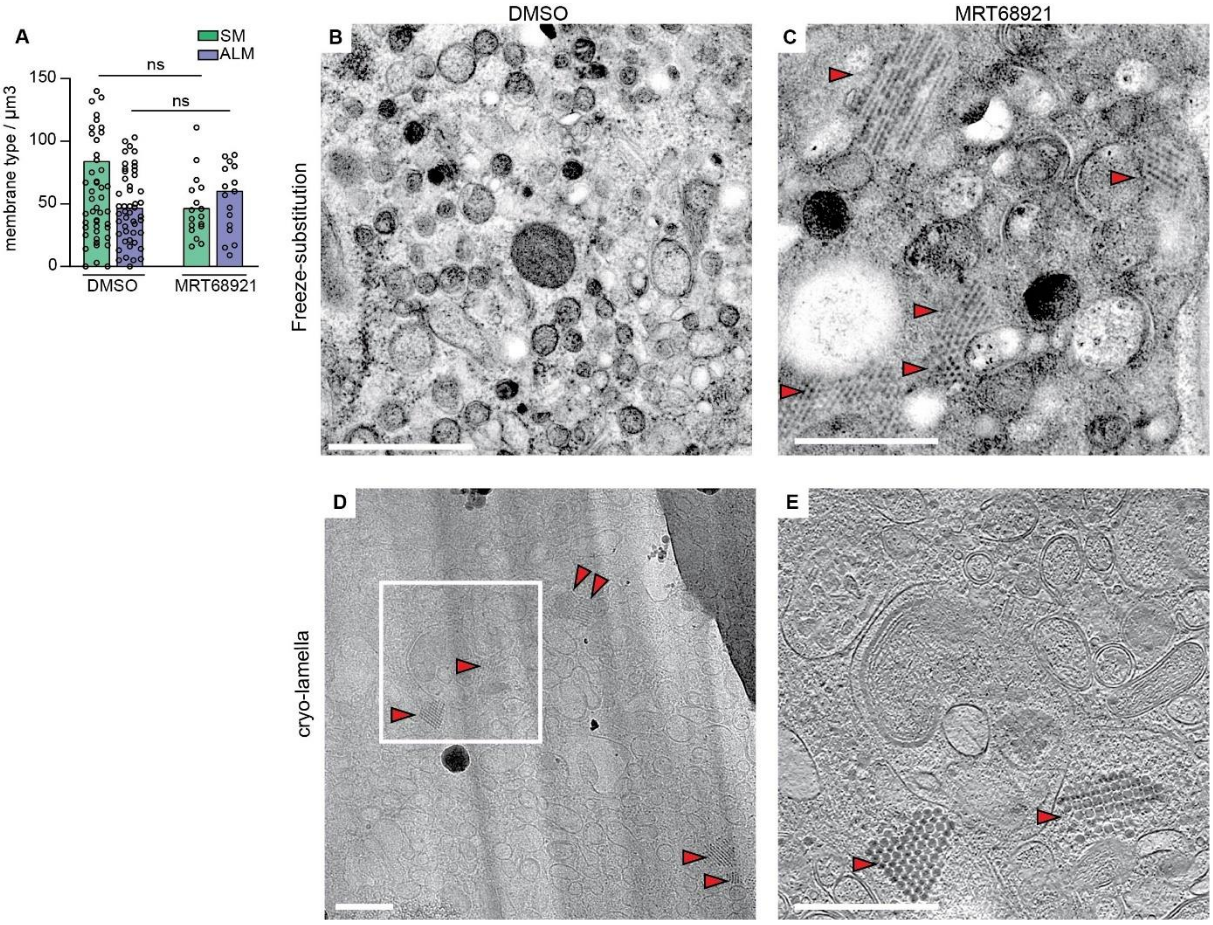


**Figure S5: Intracellular virus array formation in infected cells treated with MRT68921.** (A)Concentration of SMs vs. ALMs observed in cryo-tomograms of MRT68921 treated cells compared to untreated cells at 6 h p.i. Each dot corresponds to one tomogram analyzed and bars represent the mean (see also Supplementary table 2). (B-C) Electron micrographs of thin sections of freeze-substituted PV-infected cells in the absence (B) and presence (C) of MRT68921 at 6 h p.i., indicating the formation of virus arrays (red arrowheads) when cells are treated with MRT68921. (D) Low magnification image of cryo-lamella milled through MRT68921 treated cell at 6 h p.i. is included for comparison with (C), in which virus arrays are already identified (red arrowhead). (E) Corresponding slice through the tomogram reconstructed from tilt-series collected on the indicated region (white frames in (C)). Scale bars: 500 nm.

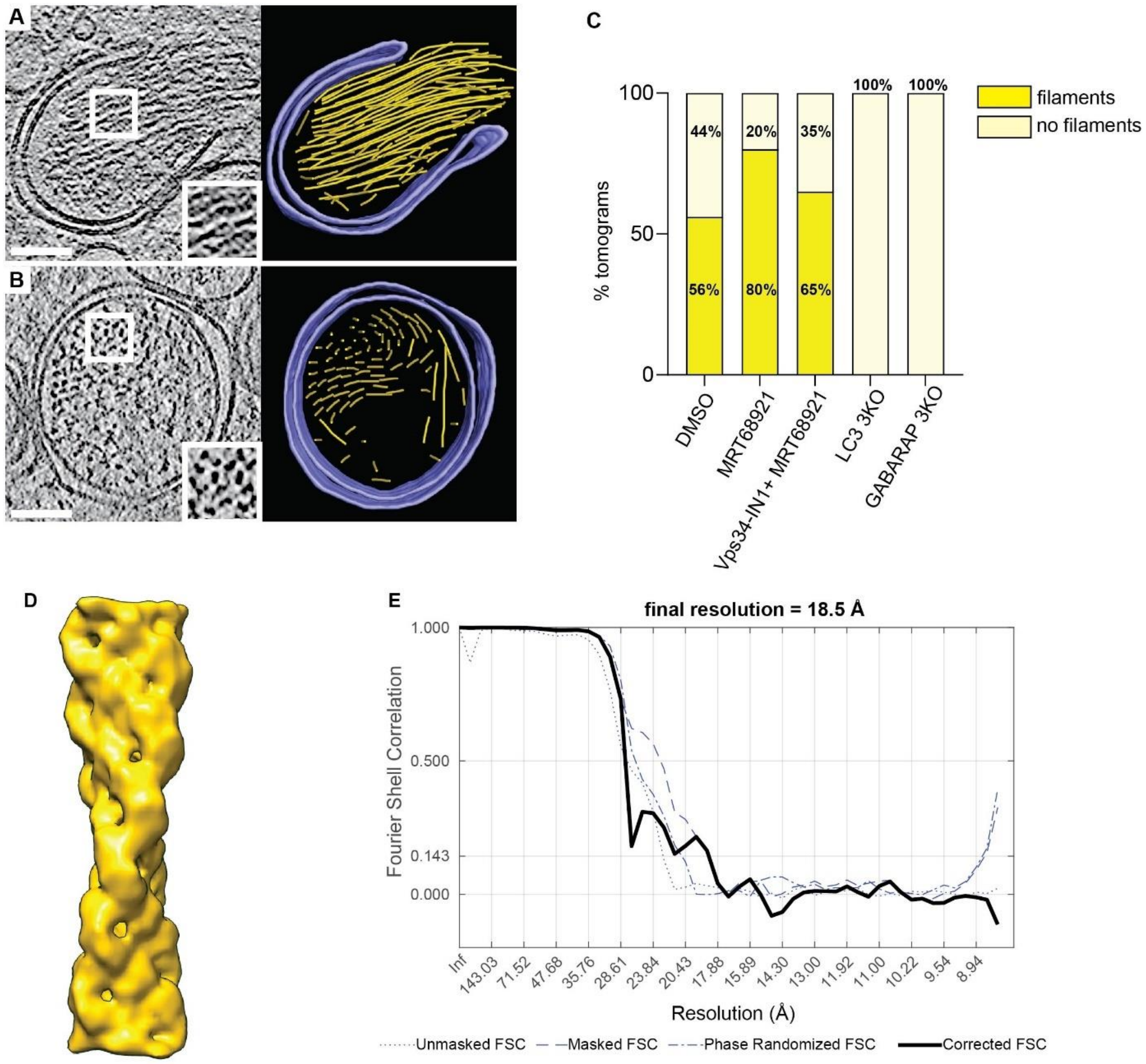


**Figure S6: ALM-associated protein filament.** (A-B) Slices through tomograms of PV-infected cells at 6 h p.i and their corresponding 3D segmentation showing large bundles of the filament (yellow) filling the interior of a phagophore-like structure (A) and double-membrane vesicle (B) (purple). Zoomed regions (white boxes) highlight the helical twist (A) and bundle formation (B) of the filaments. Scale bars: 100 nm. (C) Percentage of tomograms containing ALM-associated protein filament bundles in DMSO treated WT cells (N=34), in MRT68921 treated WT cells (N=10) and in MRT68921 + Vps34-IN1 treated WT cells (N=20), compared to LC3 (N=17) and GABARAP (N=13) 3KO cells. (D) Subtomogram average of the filaments at 18.5 Å resolution. (E) Fourier shell correlation curves for unmasked, masked, and phase-randomized (beyond 31 Å) half-sets. The corrected curve, equalling 0.143 at 18.5 Å resolution, is based on Chen *et al*[55] as implemented in subTOM.

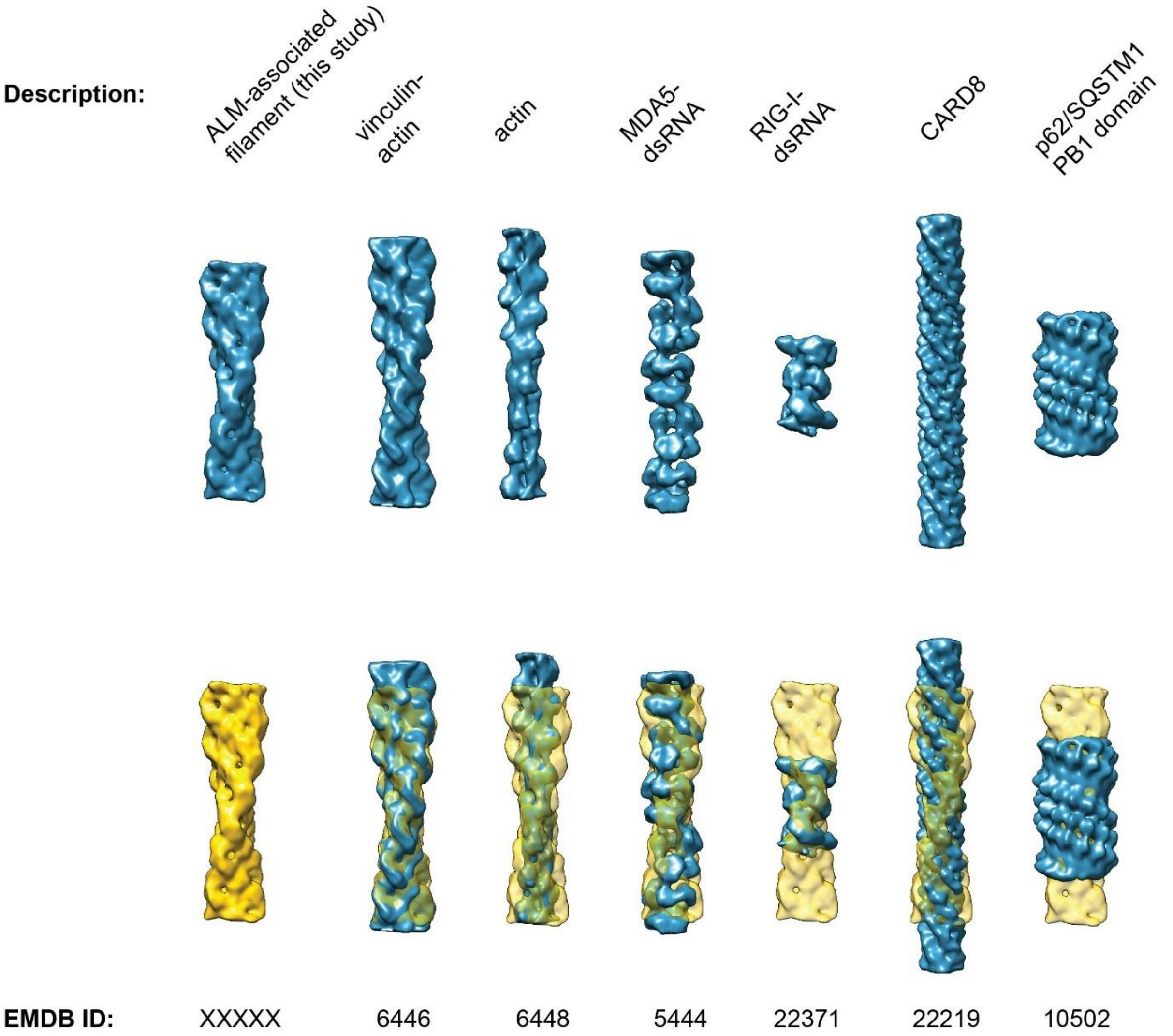


**Figure S7: Comparison of the ALM-associated filament with a selection of known filament structures.** Top row: isosurface representations of the subtomogram average of the ALM-associated protein filament from this study, and one representative of each of several classes of cellular protein filaments with known structure. In the lower row, the ALM-associated filament is shown in yellow, and semitransparent yellow when fitted to each of the other filaments using UCSF Chimera's Fit in Map function[41]. Prior to comparison, all filaments were resampled to the same voxel size and filtered to 19 Å resolution. EMDB identifiers are indicated below each volume.

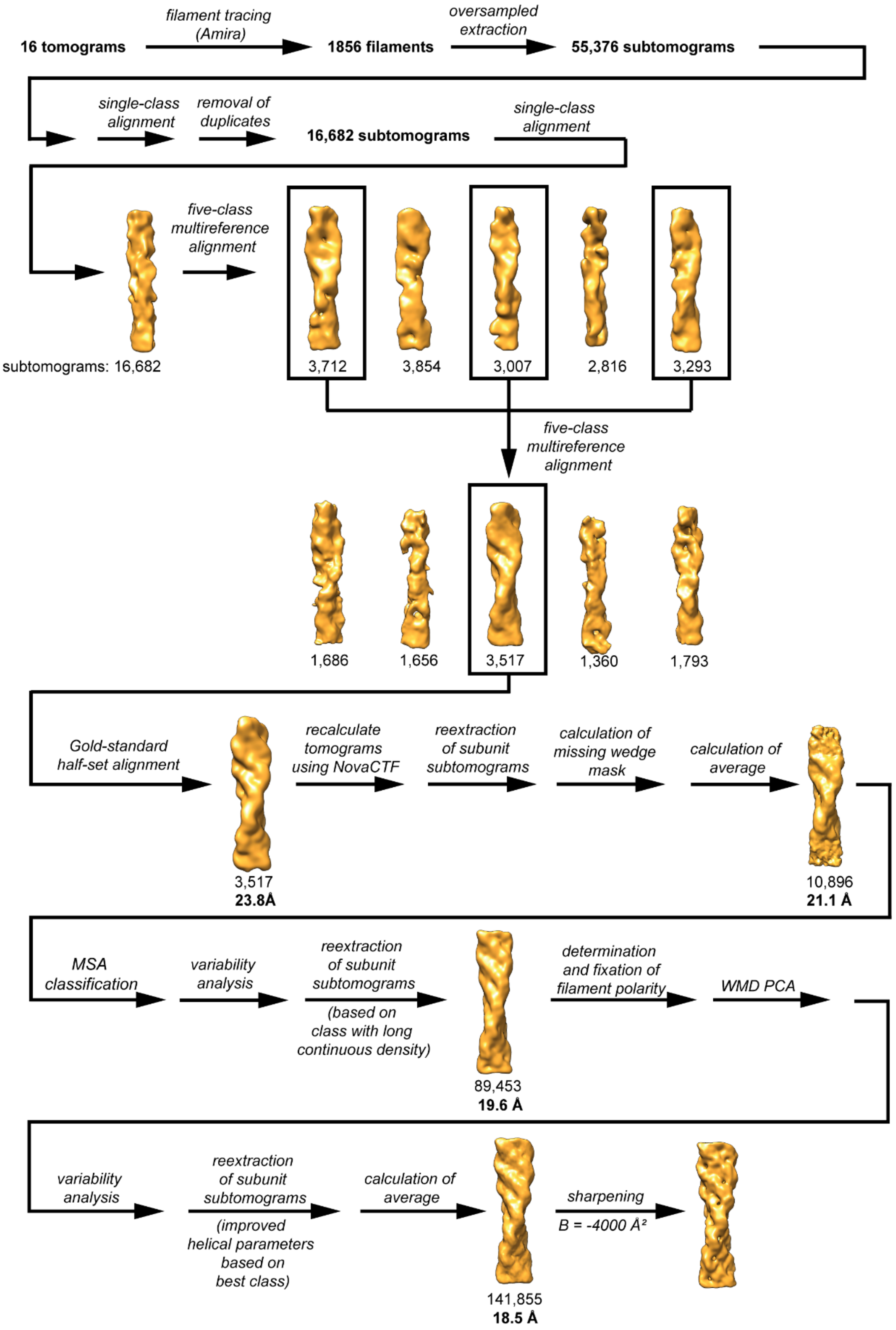


**Figure S8: Schematic of the workflow for subtomogram averaging of the ALM-associated filament.** The schematic illustrates the major steps of the data processing starting at filament tracing in the tomograms, and extraction of subtomograms along the filament axis, through the several steps of classification, reextraction and averaging.

**Table S1**: Data collection and parameter for Cryo-electron tomography of poliovirus replication and assembly sites.

| Data collection | |
|---|---|
| microscope | Titan Krios G2 |
| Voltage (KeV) | 300 |
| Camera | Gatan K2 |
| Magnification | 33,000 |
| Energy filter | yes, BioQuantum |
| Slit width (eV) | 20 |
| Pixel size in super-resolution mode (Å) | 2.18 |
| Defocus range (µm) | -3 to -5 |
| Tilt range (°) | ± 50 to ± 60 |
| Total dose (e−/Å$^2$) | 100 to 130 |
| Tomograms acquired | 168 |

**Table S2**: Concentration of membrane structures and virions (Mean ± SD).

| | N tomograms | SM / µm$^3$ | ALM / µm$^3$ | RNA-loaded virions / µm$^3$ | Empty Capsids / µm$^3$ |
|---|---|---|---|---|---|
| PV 6 hpi | 51 | 70,04 ± 58,08 | 1,21 ± 1,31 | 10,11 ± 14,74 | 6,26 ± 9,05 |
| PV 3 hpi | 14 | 84,12 ± 92,66 | 45,08 ± 29,23 | 106,11 ± 87,66 | 40,47 ± 48,91 |
| Hydantoin | 17 | 48,11 ± 25 | 60,47 ± 38,56 | 546,23 ± 737,61 | 127,53 ± 114,82 |
| MRT68921 | 19 | 21,42 ± 27,85 | 17,45 ± 20,69 | 2,21 ± 3,12 | 3,78 ± 5,3 |
| Vps34-N1+MRT68921 | 31 | 68,16 ± 44,88 | 29,17 ± 49,64 | 50,55 ± 54,84 | 152,83 ± 88,98 |
| 3KO LC3 | 23 | 54,52 ± 47,52 | 28,01 ± 27,21 | 39,61 ± 112,5 | 19,19 ± 40,92 |
| 3KO GABARAP | 13 | 53,69 ± 28,1 | 46,37 ± 31,47 | 83,01 ± 84,24 | 32,94 ± 42,73 |

**Table S3: Protein filament structures from the electron microscopy data base compared to the ALM-associated filament.** The filaments are listed by type. Fig. S7 shows the subset of filaments being most relevant either by biological function or close structural match.

| **class of filament** | **EMDB accession codes (EMD-)** |
|---|---|
| F-actin | 11976, 6448 |
| decorated F-actin | 4346, 6446, 7831, 20711, 20843, 20844, 21155, 21925, 30085 |
| CARD domain | 6842, 7314, 8902, 8903, 9332, 9943, 9948, 22219, 22220 |
| caspase | 8300 |
| CTP synthase | 0840, 8474 |
| DMC1 | 30311 |
| Dvl2/DIX | 21148 |
| glucokinase | 20309 |
| IMPDH | 4402, 8690 |
| NLRP6 | 0438 |
| MDA5-dsRNA | 0143, 4338, 4341, 5444 |
| MxB | 8577 |
| MyD88 | 4405 |
| p62/SQSTM1 | 10499, 10500, 10501 10502 |
| phosphofructokinase | 8542 |
| RAD51 | 8183, 9566 |
| RIG-I-dsRNA | 22371 |
| Torsin | 20076 |
| VPS24 | 11212 |
